# Nuclear pores safeguard the integrity of the nuclear envelope

**DOI:** 10.1101/2024.02.05.578890

**Authors:** Reiya Taniguchi, Clarisse Orniacki, Jan Philipp Kreysing, Vojtech Zila, Christian E. Zimmerli, Stefanie Böhm, Beata Turoňová, Hans-Georg Kräusslich, Valérie Doye, Martin Beck

## Abstract

Nuclear pore complexes (NPCs) constitute giant channels within the nuclear envelope that mediate nucleocytoplasmic exchange. NPC diameter is thought to be regulated by nuclear envelope tension, but how such diameter changes are physiologically linked to cell differentiation, where mechanical properties of nuclei are remodeled and nuclear mechanosensing occurs, remains unstudied. Here we used cryo-electron tomography to show that NPCs dilate during differentiation of mouse embryonic stem cells into neural progenitors. In Nup133-deficient cells, which are known to display impaired neural differentiation, NPCs however fail to dilate. By analyzing the architectures of individual NPCs with template matching, we revealed that the Nup133-deficient NPCs are structurally heterogeneous and frequently disintegrate, resulting in the formation of large nuclear envelope openings. We propose that the elasticity of the NPC scaffold mechanically safeguards the nuclear envelope. Our studies provide a molecular explanation for how genetic perturbation of scaffolding components of macromolecular complexes causes tissue-specific phenotypes.

## Introduction

The nucleus is the eukaryotic organelle that stores genetic material, and its secure maintenance is essential for cell survival. The nucleus is surrounded by the nuclear envelope, consisting of two lipid bilayers, the outer and inner membranes. Nuclear pore complexes (NPCs) are embedded in the nuclear envelope and regulate nucleocytoplasmic exchange. Structurally, the NPC has an 8-fold symmetric, cylindrical architecture, which can be subdivided into three rings stacked along the central axis: two outer rings, cytoplasmic ring (CR) and nuclear ring (NR), at the respective side of the nuclear envelope, and the inner ring (IR) at the fusion point of the outer and inner nuclear membranes.^1,2^ Each ring is composed of specific sub-complexes that are formed by multiple protein components called nucleoporins (Nups). In mammalian NPCs, the Y-complex (or Nup107-160 complex)^3–5^ oligomerizes into two tandem rings^6^ and forms the scaffold of the CR and NR.^7–10^ This ring formation is mediated by a head-to-tail contact between Nup133 at the tail of one Y-complex and Nup160 of the adjacent Y-complex.^9,11^ Despite the structural importance of the Y-complex for the overall NPC architecture, mutations in Y-complex Nups are known to affect the development and function only of specific tissues, such as kidney,^12–14^ ovary,^15^ or brain.^16–18^ Moreover, certain Y-complex Nups, including Nup133, are dispensable in mouse embryonic stem cells,^19–22^ while their absence severely affects cell differentiation.^19,21–23^ Such tissue and cell type-specific phenotypes caused by genetic defects in scaffolding Nups of central structural importance are difficult to conceive because they cannot be explained by global NPC misassembly or malfunction. Thus far, NPC architectures under genetic perturbation of scaffold Nups have not been well-studied, and the mechanisms underlying such tissue and cell type-specific defects remained unclear.

NPCs are known to be conformationally dynamic within cells. For instance, NPCs in *Schizosaccharomyces pombe* reversibly constrict under hyper-osmotic stress, where nuclear shrinkage occurs and nuclear envelope ruffling is observed.^24^ Similarly, human NPCs have a larger diameter *in situ*^25–27^ as compared to isolated nuclear envelopes,^9^ where mechanical forces are alleviated. Thus, it has been proposed that nuclear envelope membrane tension regulates NPC diameter.^24^ Intriguingly, this NPC diameter change is suggested to be relevant to nuclear mechanosensing. The nucleus is physically linked to the cytoskeleton network,^28^ and mechanical forces sensed by the cytoskeletal filaments at the cell periphery are transmitted to the nucleus and impose mechanical stress.^29–31^ This in turn triggers nuclear mechanosensing responses, such as nuclear import of the transcription factor YAP.^32^ Nuclear mechanosensing broadly alters nucleocytoplasmic transport capacity,^33^ which is thought to be mediated by increased NPC permeability due to their tension-induced deformation.^32,33^

Nuclear mechanosensing is involved in multiple biological processes, such as cell differentiation and development. In particular, cell differentiation is known to depend on the mechanosensing of the matrix stiffness.^34^ Moreover, cell differentiation also involves remodeling of nuclear properties, including nuclear stiffening^35,36^ and cytoskeleton-mediated nuclear shaping.^37^ These previous findings highlight the importance of nuclear mechanosensing during cell differentiation, and indicate that the mechanical load on the nuclear envelope increases during differentiation. Thus, it is conceivable that NPC architecture is also affected during this process, but this possibility has not been examined to date.

To address if there is a link between the NPC diameter change, cell differentiation, and perturbation of the NPC scaffold architecture, we used cryo-electron tomography and structurally analyzed NPC architecture during embryonic stem cell differentiation in wild-type and Nup133-deficient cell lines. Specifically, we hypothesized that the NPC scaffold conformationally responds to the changing mechanical environment of the nuclear envelope during differentiation, and that such a response may be impaired when the NPC scaffold architecture is perturbed. We thus compared the architectures of the NPC between these two cell lines and between the pluripotent and differentiated neural progenitor states.

## Results

### Nuclear pores dilate during early neuronal differentiation

To investigate possible changes in NPC architecture during cell differentiation, mouse embryonic stem cells (mES) were differentiated into neural progenitor cells as previously described (Fig. S1A-C).^38^ Consistent with previous reports, most of neural progenitor cells expressed the respective marker Pax6 (Fig. S1D), indicating successful differentiation. To improve the morphological consistency of vitrified samples, mES cells were synchronized to early S-phase, where the nuclear size and DNA content are more homogeneous. Clusters of cells were dissociated by trypsinization, pipetted onto EM grids, plunge-frozen and subjected to specimen thinning by cryo-focused ion beam (cryo-FIB) milling (Fig. S1E-J). We acquired 260 and 176 cryo-electron tomograms of the wild-type mES and neural progenitor cells, respectively, and structurally analyzed their NPCs by subtomogram averaging (STA) (Table S1, see methods for detail).

The subtomogram averages of the NPC from wild-type mES cells were resolved to ∼30 Å resolution (Fig. 1A, B, S2). As expected from the close phylogenetic relationship between human and mouse, the overall architecture of the wild-type mES NPC is highly reminiscent to that of the human NPC, including the density of Nup133 (Fig. S3). In addition to the structural features previously resolved in human NPC maps, a protrusion from the NR, likely corresponding to nuclear basket filaments, is resolved at the Nup107/Nup133 region (Fig. 1A, B, S3). The cryo-EM maps of the NPC from differentiated neural progenitor cells also showed almost identical structures to those of NPCs in mES cells (Fig. 1C, S2C), indicating that the subunit composition of the NPC remains largely similar during early neuronal differentiation, at least up to the neural progenitor state. However, in comparison to the NPC from the mES cells, the cryo-EM map of the NPC from the neural progenitor cells shows an inward movement of the CR and higher nuclear envelope curvature at the outer and inner nuclear membrane fusion point (Fig. 1C), both of which are indicative of NPC dilation.^24,27^ Measurements of individual NPC diameters based on the subtomogram averages of opposing subunits (see methods for details) indeed confirmed that wild-type neural progenitor NPCs have a significantly larger diameter than wild-type mES NPCs (Fig. 1D), thus supporting our notion that NPC architecture is affected during cell differentiation.

**Figure 1:**
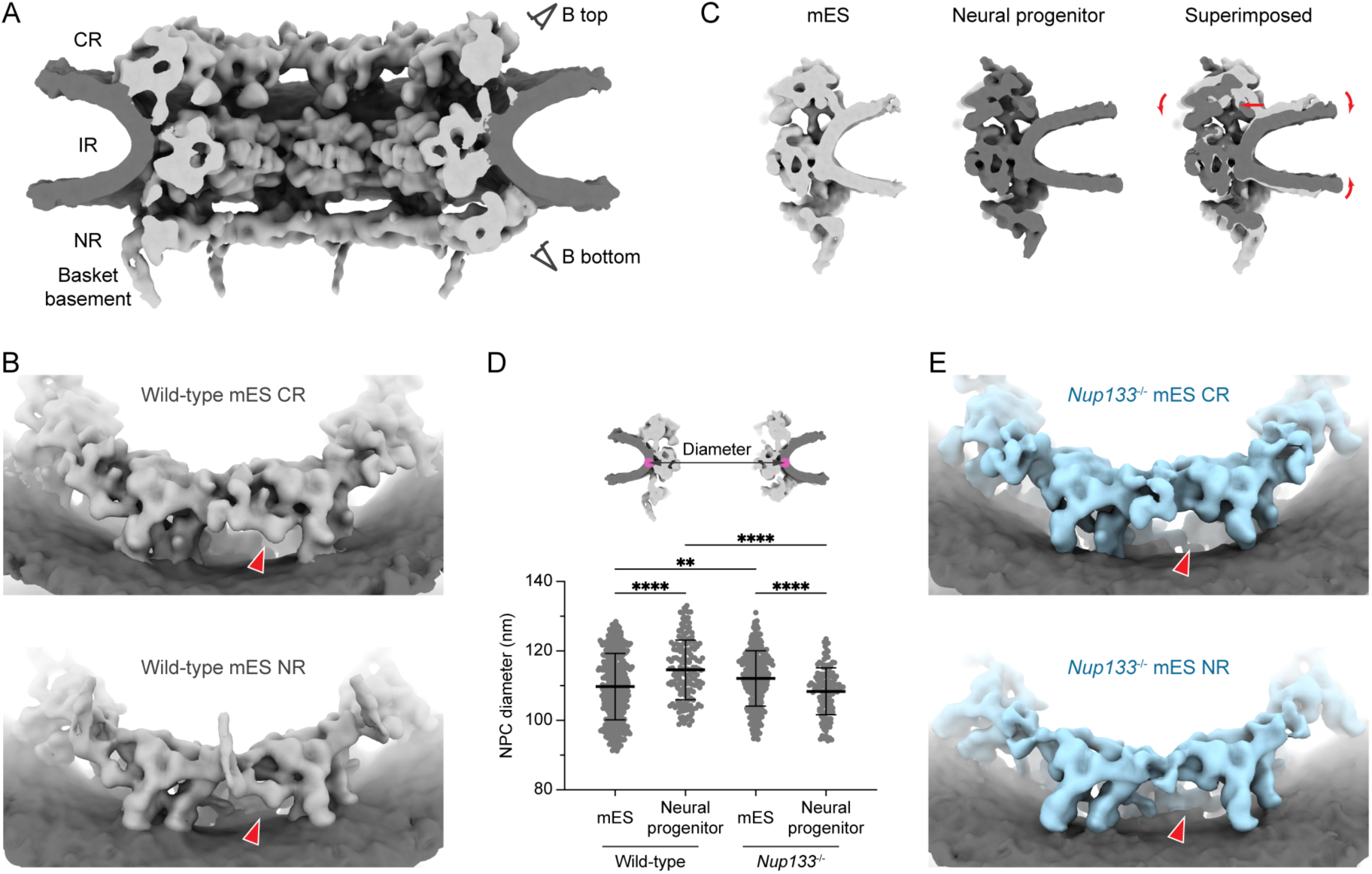
Architectures of mES and neural progenitor NPCs. (A) Composite cryo-EM map of the wild-type mES NPC shown as a cutaway view. Viewing angles of panel (B) are indicated by eye symbols. CR; cytoplasmic ring, IR; inner ring, NR; nuclear ring. (B) Cryo-EM maps of the CR (top) and NR (bottom) of the wild-type mES NPC. (C) Structural comparison of the wild-type mES NPC and the wild-type neural progenitor NPC. Single subunit is shown as a cutaway side view. Two maps are superimposed based on the position of the IR protomer, and the relative shift of the CR and nuclear membrane in the neural progenitor NPC map in comparison to the mES NPC map is indicated by red arrows. (D) NPC diameter measurements based on subtomogram averages. Schematic on top illustrates the measured distance between two opposing IR protomers. Graph (lower panel) depicts the mean values of measured diameters with standard deviations shown as black bars (n = 446 NPCs for the wild-type mES cells, n = 173 NPCs for the wild-type neural progenitor cells, n = 317 NPCs for the *Nup133*^-/-^ mES cells, n = 138 NPCs for the *Nup133*^-/-^ neural progenitor cells). Kruskal-Wallis rank sum test; **p < 0.01, ****p < 0.0001. (E) Cryo-EM maps of the CR (top) and NR (bottom) of the *Nup133*^-/-^ mES NPC, shown as in (B). In (B) and (E), the positions of Nup133 are highlighted with red arrowheads.

### Nuclear pores devoid of Nup133 retain remnant contacts between neighboring protomers

Nup133 is known to mediate the head-to-tail connection between adjacent Y-complexes and to bridge neighboring protomers of the CR and NR.^9,11^ Despite this critical scaffolding role^5^ and its clear presence within the averages of the wild-type NPC (Fig. 1B), Nup133 is dispensable for the proliferation of mES cells.^19,20,22^ In Nup133-deficient cells, however, its absence is detrimental for neuronal differentiation, and results in reduced growth and increased cell death upon differentiation induction.^19,22^ This differential requirement for Nup133 at distinct cell differentiation states challenges our present structural understanding of NPC architecture, which would have predicted a strict requirement for Nup133 in maintaining the NPC scaffold. We therefore reasoned that mES cells devoid of Nup133 would be a suitable model system to further assess the importance of the Y-complex for the structural integrity of NPC architecture during differentiation.

We structurally analyzed NPCs in previously generated *Nup133*^-/-^ mES cells.^20^ As expected, the cryo-EM maps of the *Nup133*^-/-^ mES NPC lack the density for Nup133 in the CR and NR (compare Fig. 1B to E), while the IR shows a structure almost identical to that of the wild-type mES NPC (Fig. S2B-D). Strikingly, even without Nup133, the CR and NR appear to have an overall intact structure. A connection between the adjacent protomers is observed in the Nup107 - Nup205/Nup93 heterodimer region in a consistent manner to that in the wild-type mES NPC (Fig. 2), highlighting a previously underappreciated importance of this additional head-to-tail contact interface. This interaction between two adjacent protomers could be particularly important for the NR, since additional components that would support structural integrity, such as cytoplasm-specific Nup358, are absent. In accordance with the previously reported nuclear basket misassembly and increased dynamics of a basket component Nup in the *Nup133*^-/-^ cells,^20^ the respective protrusion in the Nup107/Nup133 region was diminished (compare Fig. 1B to E, Fig. 2).

**Figure 2:**
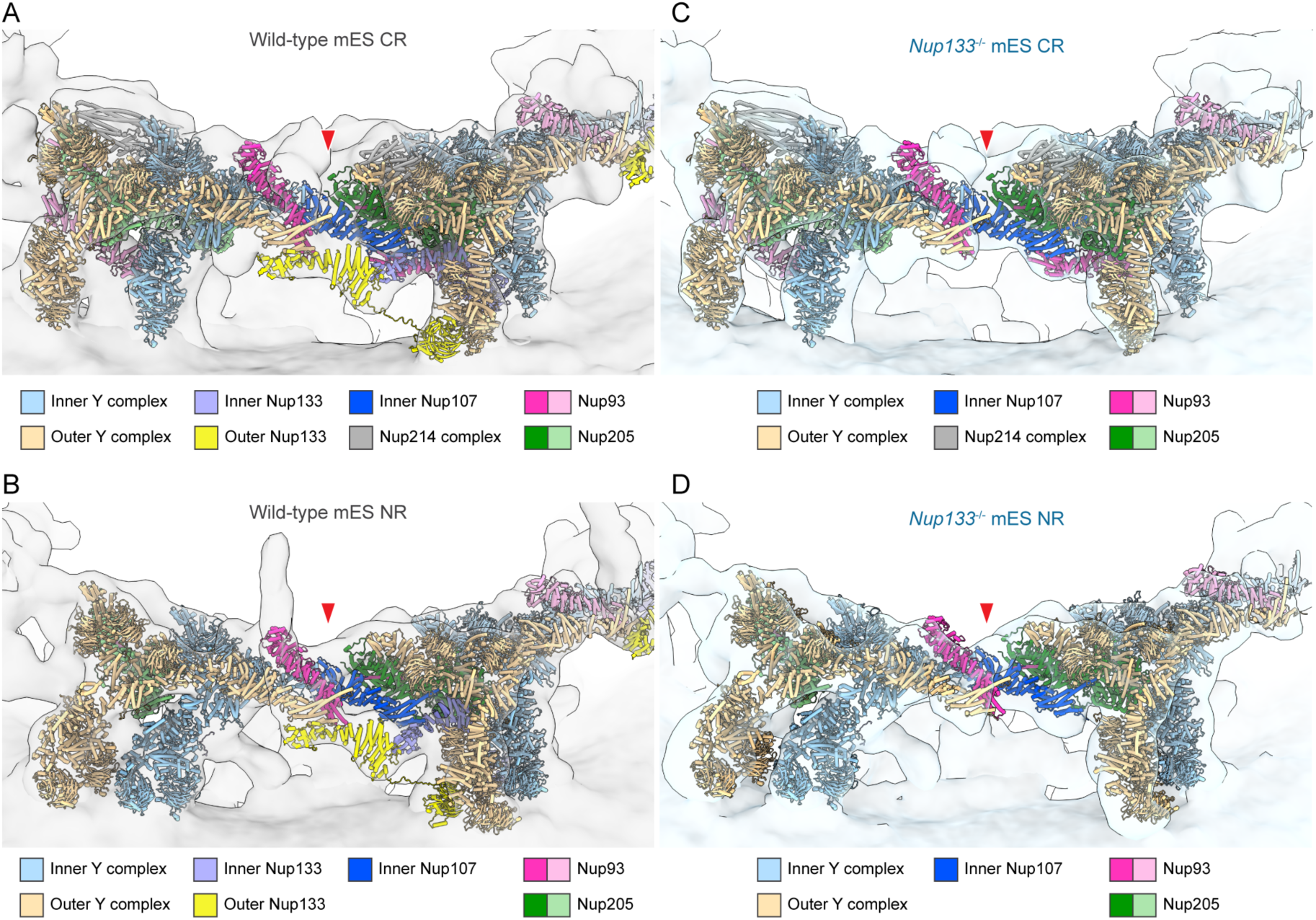
The density for the head-to-tail contact is present in the *Nup133*^-/-^ mES NPC. (A - D) The head-to-tail contact between the CR (A) and NR (B) protomers of the wild-type mES NPC, and the CR (C) and NR (D) protomers of the *Nup133*^-/-^ mES NPC. The CR and NR models of dilated human NPC (PDBID: 7R5J) are fitted into the corresponding cryo-EM maps. Nup93 and Nup205 molecules involved in the head-to-tail contact at the center of the images are highlighted in bold color. The main interaction interfaces are highlighted with red arrowheads. In (A) and (C), the model of Nup358 is omitted for clarity of the figure.

### Nuclear pores devoid of Nup133 constrict during early neuronal differentiation

Nup133-deficient cells are able to differentiate into neural progenitor cells, albeit with abnormally maintained characteristics of the pluripotent state,^19^ and ultimately fail to terminally differentiate into postmitotic neurons.^19,22^ We therefore structurally analyzed NPC architecture in neural progenitor cells (Fig. S1D) obtained from the *Nup133*^-/-^ mES cells. Cryo-EM maps of the NPC from the *Nup133*^-/-^ neural progenitor cells were reminiscent to those of the *Nup133*^-/-^ mES NPC (Fig. S2D, E). Surprisingly however, the diameter measurements revealed a constriction of the *Nup133*^-/-^ neural progenitor NPCs in comparison to the *Nup133*^-/-^ mES NPCs (Fig. 1D). This is in contrast to the wild-type mES cells, where we observed an NPC dilation within the differentiated neural progenitor cells. Since NPCs cannot actively change their diameter and rather passively react to external mechanical forces,^24^ these data imply that the effects of differentiation on nuclear envelope properties, such as nuclear envelope tension, are improperly propagated to the NPCs in the *Nup133*^-/-^ neural progenitor cells. Overall, these findings point towards a potential link between the structural integrity of the Y-complex and the mechanical properties of nuclear envelope itself.

### A subpopulation of NPCs devoid of Nup133 has non-canonical symmetries

To understand how the loss of Nup133 in the NPC architecture is linked to the physical properties of the nuclear envelope, we further characterized NPCs within the *Nup133^-/-^* mES cells. Visual inspection of the reconstructed tomograms revealed that NPCs with non-canonical 7- or 9-fold symmetric architectures are present in the tomograms from the *Nup133*^-/-^ mES cells, in addition to the canonical 8-fold symmetric NPCs (Fig. 2A-F). This is surprising, because non-8-fold symmetric NPCs are thought to be very rare.^39^ By reference-based classification (see Methods for details), 34 and 32 out of 400 particles were classified as 7-fold and 9-fold symmetric NPCs, respectively, while 323 particles were attributed to 8-fold symmetry (Fig. S4A). The remaining 11 particles showed ambiguous class assignment and were thus discarded from further analysis. Similar classification using the particles of the wild-type mES NPC resulted in one class with 8-fold symmetric architecture (Fig. S4B). These results indicate that a considerable subpopulation of the NPCs in the *Nup133*^-/-^ mES cells has non-canonical symmetries, whereas NPCs with aberrant symmetries are rarely observed in the wild-type mES cells.

We further characterized NPCs with non-canonical symmetries using subtomogram averaging. The particle sets of 7-fold and 9-fold symmetric NPCs yielded the moderately resolved averages of the IR (Fig. S4C-D). Although the resolution is reduced due to the limited number of particles, the observed structural features are overall consistent with canonical IRs (Fig. 3G-I). In contrast, the averages of the CR and NR of 7-fold symmetric NPCs did not exhibit any interpretable structural features (Fig. S4C). Averages of 9-fold symmetric NPCs also failed to show clear NR-like architecture (Fig. S4D), while structural features typical for the CR were apparent (Fig. 3I, S4D). These observations indicate that the subpopulation of NPCs with aberrant symmetries possess intact IRs, while CRs and NRs are in part deteriorated or diminished. The observed remnant densities for CR and NR (Fig. S4C-D) may be indicative of further structural heterogeneity. However, given the low overall particle number, these cannot be further analyzed by averaging-based methods such as STA.

**Figure 3:**
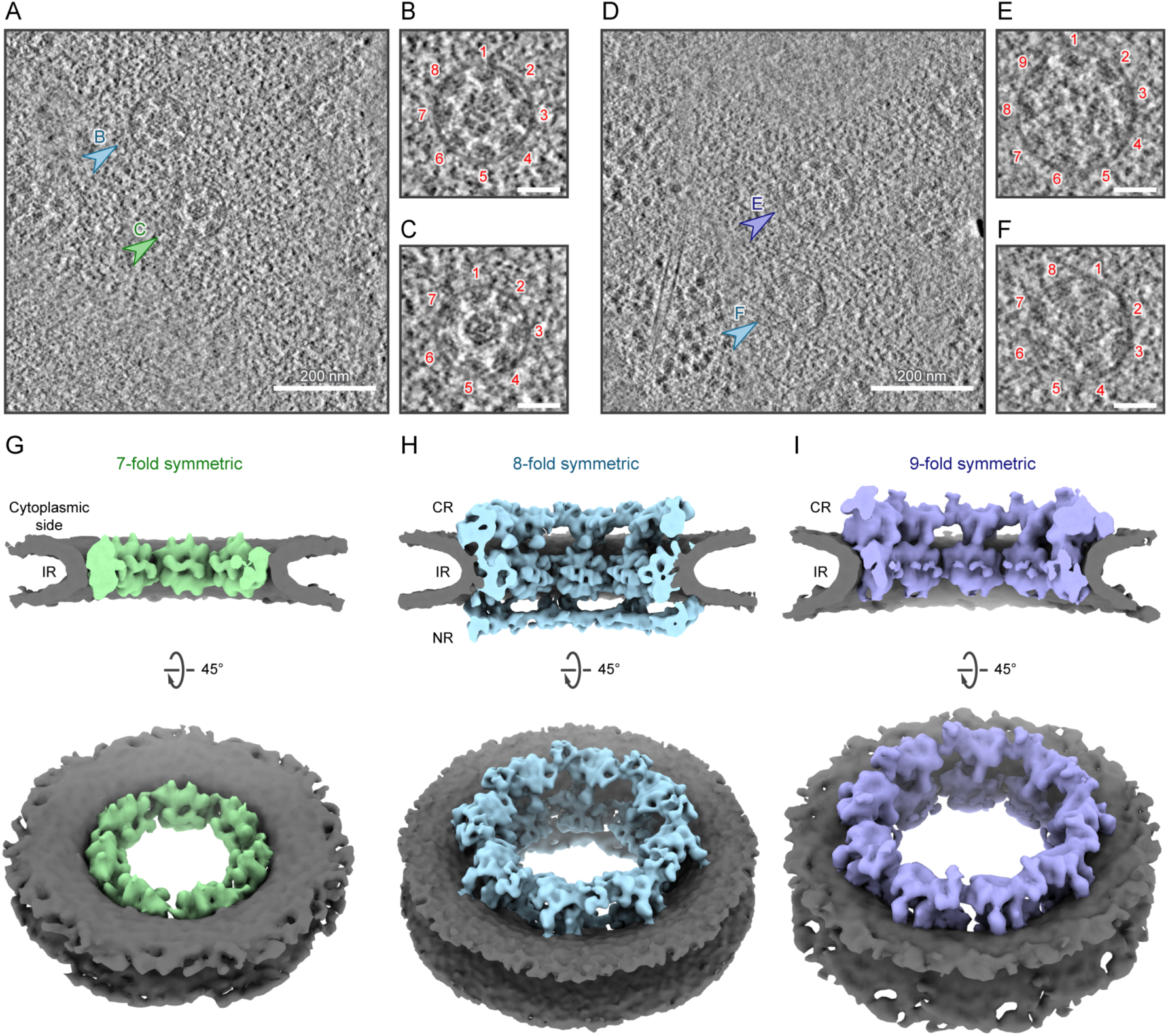
NPCs with non-canonical symmetries are present in *Nup133*^-/-^ mES cells. (A - F) Representative slices from reconstructed tomograms of the *Nup133*^-/-^ mES cells, showing top views of the 7-fold (A, C), 8-fold (A, B, D, and F) and 9-fold (D, E) symmetric NPCs. In (A) and (D), the 8-fold symmetric NPCs are indicated with light blue arrowheads, while the NPCs with 7-fold (C) and 9-fold (E) symmetric architectures are indicated with green and purple arrowheads, respectively. The top views in (A) and (D) are shown as enlarged views in (B, C, E, and F) with protomers numbered. Scale bars in (B, C, E, and F), 50 nm. (G – I) Composite cryo-EM maps of the 7-fold (G), 8-fold (H), and 9-fold (I) symmetric NPCs, shown as a cutaway view (top) and a cytoplasmic view (bottom). Note that the CR and NR of the 7-fold symmetric NPC and the NR of the 9-fold symmetric NPC did not yield interpretable subtomogram averages and thus are not included in the composite cryo-EM maps shown in (G) and (I).

### NPCs devoid of Nup133 are highly heterogenous and have an incomplete ring architecture

Three-dimensional template matching (TM)^40,41^ is a method that can be used to address the challenge of analyzing such structural heterogeneity. TM detects the structural signature of target molecules within cryo-electron tomograms by cross-correlating a reference structure with all possible locations and orientations of a given tomogram, without the need for averaging. We have recently shown that this method can detect individual NPC subunits,^42^ thus opening up the exciting possibility to examine the heterogenous presence of NPC subunits in more detail. We used the cryo-EM maps of the CR, IR and NR protomers as search templates and analyzed the ultrastructure of individual NPCs with non-canonical symmetry that were fully contained in the respective tomograms acquired from the *Nup133*^-/-^ mES cells (Fig. 4A). In these selected particles, IR protomers were detected as complete or almost complete rings with the expected 7- or 9-fold symmetries (Fig. S5A, B). However, neither CR nor NR protomer was detected in almost half of the 7-fold symmetric NPCs, indicating a complete absence of the respective rings (Fig. 4B, C). The remaining particles showed the presence of two to five CR protomers (Fig. 4C) arranged in a partially open ring-like architecture, while NR protomers were less frequently detected (Fig. 4C). In 9-fold symmetric NPCs, one third of the tested particles showed complete CRs, while the rest contained five to eight protomers arranged in an incomplete ring architecture (Fig. 4D, E). The number of detected NR protomers was more variable, yet the majority of the 9-fold symmetric NPCs had a partially open NR architecture (Fig. 4E).

**Figure 4:**
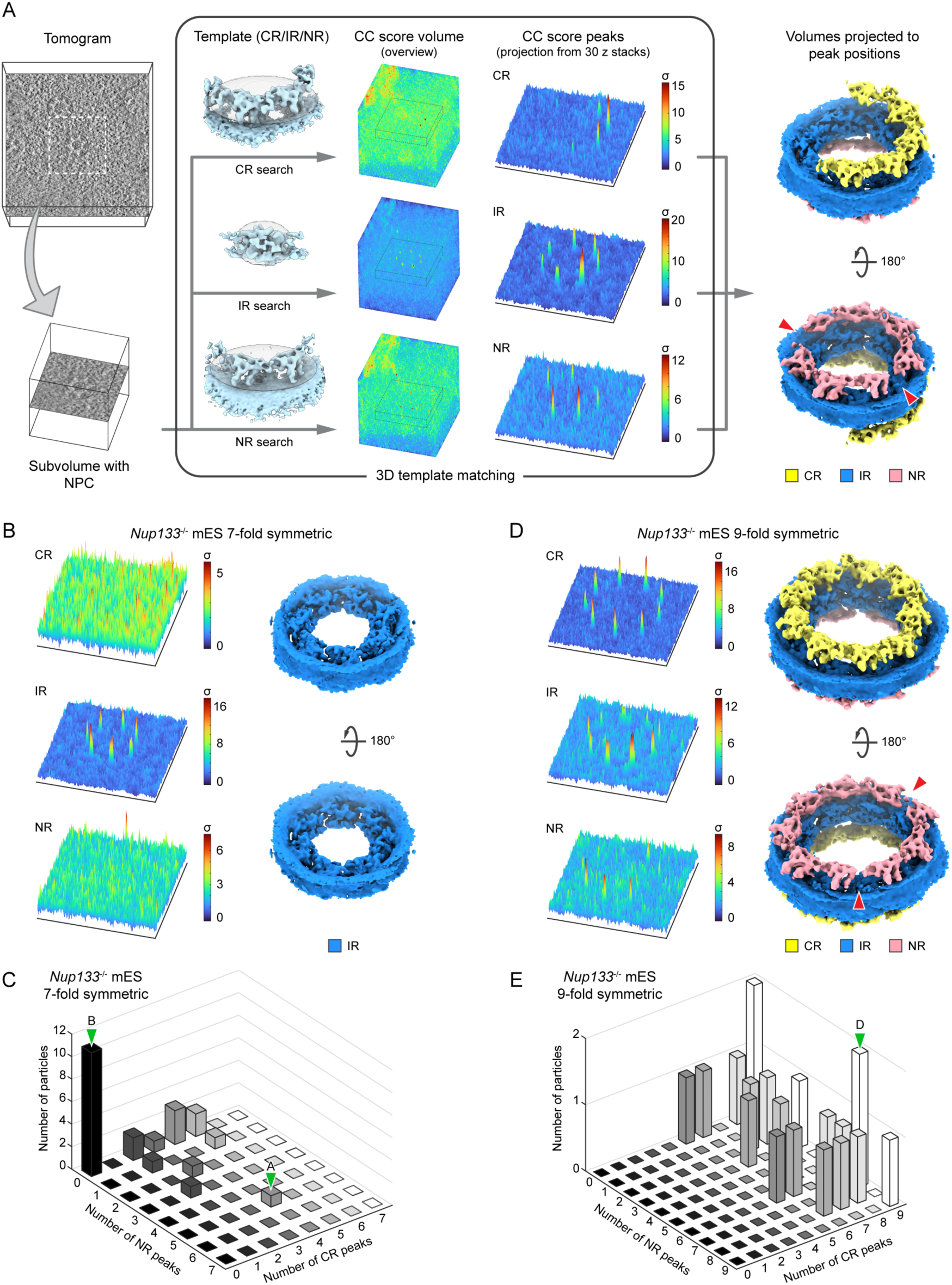
TM analysis of the 7-fold and 9-fold symmetric NPCs shows incomplete CR and NR ring architectures. (A) Schematic showing the overall workflow of the TM analysis. Subvolumes containing NPCs of interest are extracted from tomograms and subjected to three TM runs, using the CR, IR and NR averages as the search templates. Ellipsoidal masks used for the TM runs are shown as grey transparent spheres around the averages. The positions of the peaks were extracted from the output cross-correlation (CC) score volumes (see methods for details). For the visualization of the peaks, ±15 z-stacks around the cross-correlation peaks are extracted, and maximum values along z axis are presented as 3D plots to show multiple peaks with different z coordinates in a single plot. Based on the positions of the extracted peaks and corresponding template orientations, the maps of the templates are projected back to generate a pseudo-composite map (rightmost panel). (B) Representative results of the TM analysis of a 7-fold symmetric NPC. 3D plots of the peaks and a pseudo-composite map are shown as in (A). (C) 3D histogram showing the number of 7-fold symmetric NPCs and their differing numbers of CR and NR peaks detected by the TM analysis (n = 24 NPCs). (D) Representative results of the TM analysis of a 9-fold symmetric NPC. 3D plots of the peaks and a pseudo-composite map are shown as in (A). (E) Analysis as in (C) for the 9-fold symmetric NPCs with different numbers of CR and NR peaks (n = 20 NPCs). In the pseudo-composite maps in (A) and (D), gaps in the ring architectures are indicated with red arrowheads. In (C) and (E), bars that include the examples shown in (A, B and D) are indicated by green arrowheads.

Encouraged by these results, we next analyzed NPC subsets of similar size with canonical 8-fold symmetry. In the *Nup133*^-/-^ cells, only 4 out of 24 NPCs showed a complete CR and NR, while the majority of the particles still had incomplete architectures (Fig. 5A, B). This is contrasted by the analysis of the NPCs in the wild-type mES cells, in which 18 out of 29 particles have fully detectable CRs and NRs, and most of the remaining 11 have close to complete ring architectures (Fig. 5C, D). In both datasets, the IR protomers were similarly detected as complete or almost-complete 8-fold symmetric rings (Fig. S5C, D). These results very clearly show that 8-fold symmetric NPCs devoid of Nup133 more frequently exhibit heterogenous and incomplete CRs and NRs compared to the wild-type NPCs, and that the incomplete ring architectures can be observed globally among the NPCs in the *Nup133*^-/-^ cells, independent of their symmetries. For both the 8- and 9-fold symmetric NPCs, the NR shows higher heterogeneity in comparison to the CR (Fig. 4E, 5B), which may well explain the previously reported nuclear basket heterogeneity in the *Nup133*^-/-^ cells.^20^ Moreover, although the subtomogram averages of the NPCs from the *Nup133*^-/-^ cells appear largely normal, the overall number of NPCs with incomplete CRs and/or NRs likely exceeds 80% in the respective dataset. We thus conclude that the lack of Nup133 globally affects the structural integrity and completeness of CR and NR in addition to perturbing symmetry of a smaller subpopulation of NPCs. Importantly, the TM analysis revealed heterogeneity in the NPC architecture on the single protomer level, which would have been overlooked with averaging-based particle analysis.

**Figure 5:**
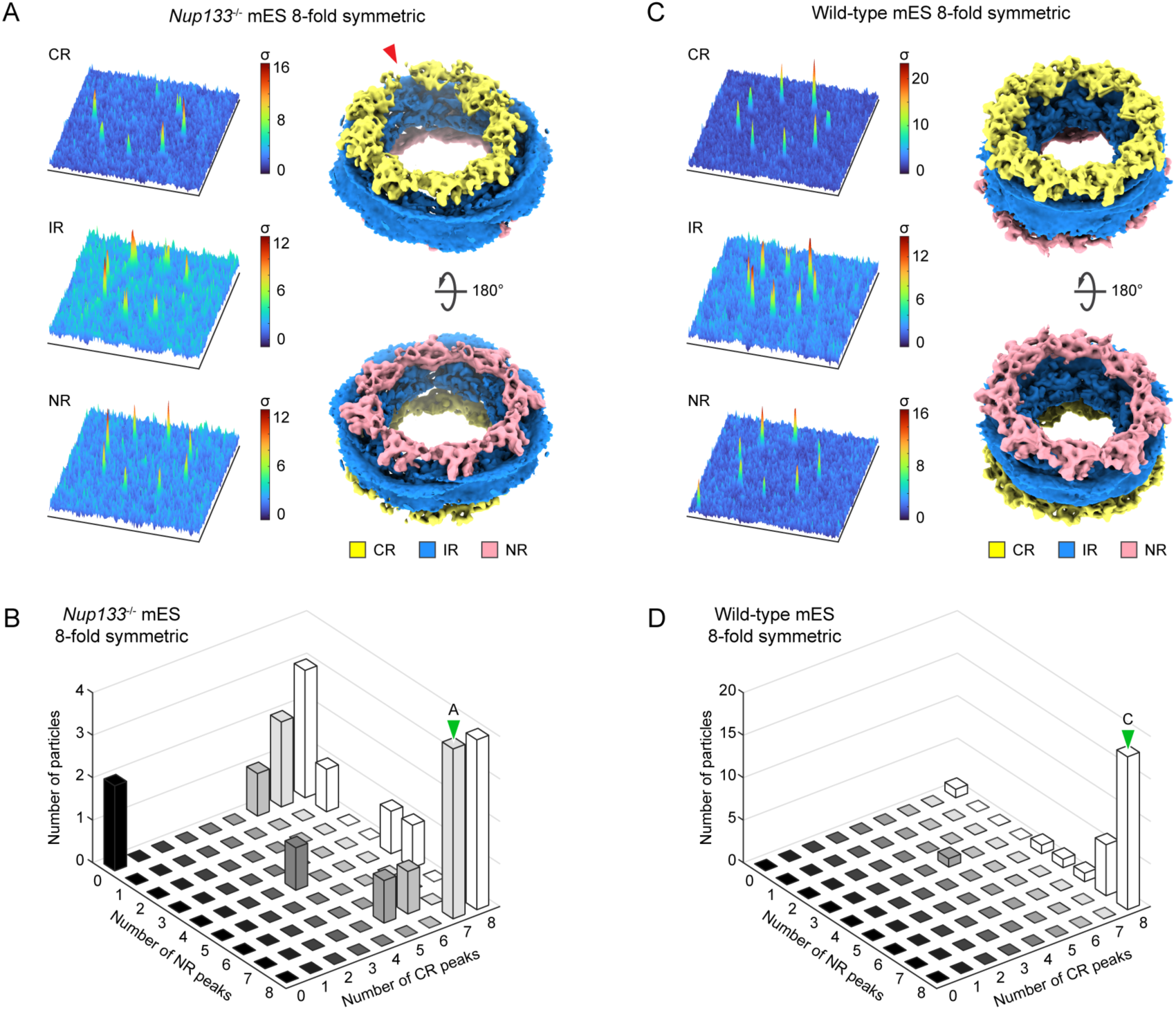
The 8-fold symmetric NPCs in the *Nup133*^-/-^ mES cells more frequently possess incomplete CR and NR architectures in comparison to the wild-type mES NPCs. (A) Representative results of the TM analysis of an 8-fold symmetric NPC in the *Nup133*^-/-^ mES cells. In the pseudo-composite map, a gap in the ring architecture is indicated with a red arrowhead. (B) 3D histogram showing the number of 8-fold symmetric *Nup133*^-/-^ mES NPCs and their differing numbers of CR and NR peaks (n = 24 NPCs). (C) Representative results of the TM analysis of an 8-fold symmetric NPC in the wild-type mES cells. (D) Analysis as in (B) for the 8-fold symmetric wild-type mES NPCs (n = 29 NPCs). In (A) and (C), 3D plots of the peaks and a pseudo-composite map are shown as in Figure 4A. In the pseudo-composite map in (A), a gap in the ring architecture is indicated with a red arrowhead. In (B) and (D), bars that include the examples shown in (A) and (C) are indicated by green arrowheads.

### NPC over-stretching results in large openings in the nuclear envelope

It has been proposed that nuclear envelope tension regulates NPC dilation, while in the absence of nuclear envelope tension, NPCs constrict into a conformational ground state.^24^ In such a scenario, one may conceptualize the NPC scaffold as an annular spring that counteracts laterally applied forces and thereby maintains the pore size within a certain range. The CR and NR architectures remain largely unaltered during NPC dilation,^10,27^ and they thus may be regarded as the source of restoring force. We therefore reasoned that NPCs with an incomplete ring architecture may have at least partially lost the spring-like properties, and further hypothesized that they may excessively dilate (over-stretch) and possibly disintegrate during differentiation due to insufficient structural support to withstand the mechanical stress. With a certain subpopulation of NPCs that are over-stretched, nuclear envelope membrane tension could be overall relieved, and this globally reduced tension would allow the majority of NPCs to constrict (Fig. 6A). Thus, our model would also explain the NPC constriction we observed in differentiated *Nup133*^-/-^ cells (Fig. 1D). This model leads to several predictions: i) large openings in the nuclear envelope should represent over-stretched NPCs and thus may contain NPC scaffolds or parts thereof; ii) over-stretched NPCs should occur more frequently upon differentiation in comparison to the pluripotent state; and iii) over-stretched NPCs should occur more frequently in the *Nup133*^-/-^ cells than in the wild-type cells.

**Figure 6:**
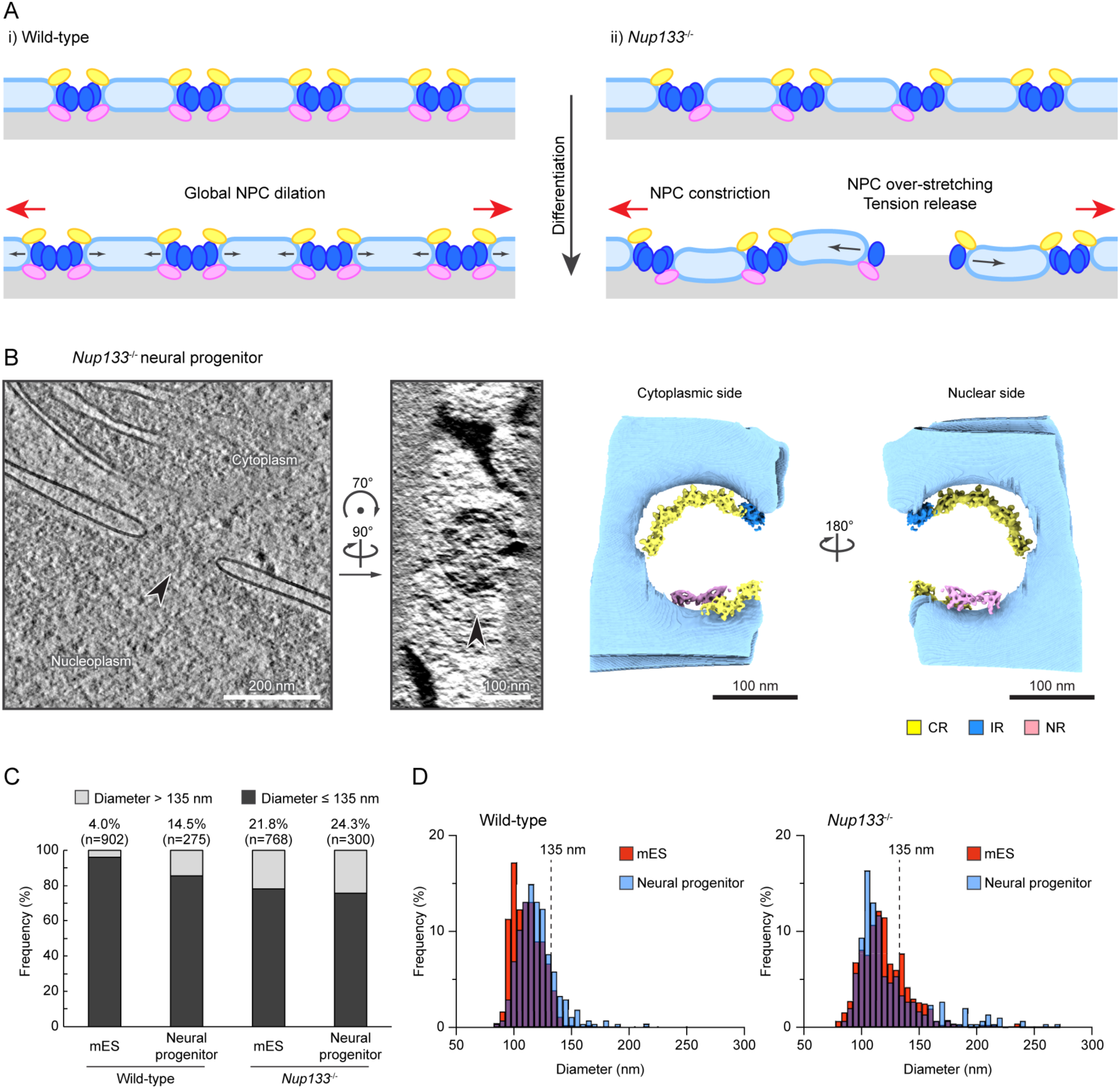
Over-stretched NPCs are present in the *Nup133*^-/-^ neural progenitor cells. (A) Hypothetical model of NPC diameter changes during neural differentiation. In the wild-type cells, the increased nuclear envelope tension during neural differentiation is homogeneously propagated along the nuclear envelope, leading to the overall dilation of the NPCs (i). In the *Nup133*^-/-^ cells, the increased nuclear envelope tension causes over-stretching of NPCs with impaired structural robustness, leading to the release of the nuclear envelope tension and the overall constriction of the NPCs (ii). Nuclear envelope is colored in light blue, and the CR, IR, and NR of the NPC are colored in yellow, blue, and pink, respectively. Red and black arrows depict nuclear envelope membrane tension and the motion of NPC scaffolds. (B) Representative example of the over-stretched NPCs observed in the *Nup133*^-/-^ neural progenitor dataset. Slices from the reconstructed tomograms showing the side views (left) and the top views (middle) of the over-stretched NPCs are presented, together with the pseudo-composite maps generated from the results of the TM analysis (right). The CR, IR, and NR in the pseudo-composite maps are colored as in Figure 3A and shown together with the segmented membrane colored in light blue. (C) Quantification of the over-stretched NPCs in the four datasets (wild-type mES cells and neural progenitor cells; *Nup133*^-/-^ mES cells and neural progenitor cells). The number of analyzed NPCs and the percentage of the over-stretched NPCs (NPCs with diameter larger than 135 nm) are indicated on top of each bar. (D) Histogram showing the distribution of the measured NPC diameters depicted in (C). Note that the overall diameter distribution is consistent to the results of subtomogram average-based measurement shown in Figure 1C.

To test the first prediction that incomplete NPCs over-stretch and create large openings in the nuclear envelope upon an increase in membrane tension, we examined the tomograms of neural progenitor cells obtained from the *Nup133*^-/-^ cells, where the aberrant NPCs with an incomplete architecture would likely have been exposed to the increased nuclear envelope tension during the induced cell differentiation. Indeed, we frequently found abnormally large holes in the nuclear envelope with diameters ranging from 150 nm to 200 nm (Fig. 6B, S6A-B), which are inconsistent with present structural models of intact NPC architecture. In 19 out of the 24 analyzed large membrane holes, TM analysis detected subunits of at least one of the three rings (Fig. S6A), confirming that the observed membrane holes are indeed lined with remnants of NPCs, presumably over-stretched and already largely disintegrated. At these sites, incomplete CRs are often detected at a position distant from the nuclear envelope (Fig. 6B, S6A), implying that they have detached from the membrane. Such membrane dissociation is less prominently observed for the NRs (Fig. S6A), possibly reflecting a stronger membrane association supported by the nuclear ring-specific component ELYS^10^ in comparison to the CRs. In addition, the IRs in these aberrant NPCs are always incomplete and individual IR protomers are often distantly spaced (Fig. S6A), which is inconsistent with an intact linker connection between two adjacent IR protomers. Taken together, these structural features indicate that the observed membrane holes are NPCs that have lost their intact architecture and have disintegrated, likely due to over-stretching.

### NPCs with compromised structural integrity more frequently disintegrate upon induced differentiation

We next examined our prediction ii) and asked whether the over-stretched NPCs were more frequently observed in the neural progenitor cells. Since the measured NPC diameters based on our subtomogram averages are distributed between 90 nm and 130 nm (Fig. 1D), we set an arbitrary threshold for the diameter of intact NPCs to a maximum of 135 nm, whereby NPCs larger than 135 nm were considered to be over-stretched. As expected, the frequency of over-stretched NPCs was elevated in the wild-type neural progenitor cells in comparison to the wild-type mES cells (Fig. 6C), consistent with the observation that NPCs dilate during differentiation and the notion that this dilation is driven by increased membrane tension. To address our last prediction, we examined the *Nup133*^-/-^ cells, in which the frequency of over-stretched NPCs is indeed drastically increased in comparison to the wild-type cells and reaches 20% (Fig. 6C). Although this frequency remains similar in neural progenitor cells, the distribution of the NPC diameter shows that the size of the over-stretched NPCs generally becomes considerably larger and more variable in neural progenitor dataset (Fig. 6D), indicating that the NPC architecture is more adversely affected upon differentiation. Together, these observations are consistent with the above-proposed model (Fig. 6A), and we conclude that an increase in nuclear envelope tension leads to NPC over-stretching, which can result in disintegration of NPCs. Although the exact causal relationships await further investigation, the frequent NPC disintegration observed within the *Nup133^-/-^* cells could well be a major contributing factor to the previously described differentiation defect phenotype.

### Cells with incomplete NPC scaffolds accumulate DNA damage during cell differentiation

In cells, NPC disintegration should affect nucleocytoplasmic compartmentalization and thus should coincide with DNA damage, which is known to increase when nuclear integrity is compromised.^43–46^ To investigate this possibility, we analyzed the presence of DNA damage in the neural progenitor cells undergoing differentiation. By quantifying the foci of phosphorylated histone H2AX (γ-H2AX), a widely used marker for DNA damage,^47,48^ we indeed found an increase of damage in the *Nup133*^-/-^ neural progenitor cells compared to the wild-type cells 24 h after induction of differentiation (Fig. 7A, B). Prior studies in human cells have shown that mechanical stress, such as osmotically induced cell swelling^49,50^ or mechanical cell stretching,^51^ alone did not induce the accumulation of γ-H2AX foci, while perturbation of the nuclear envelope properties in such mechanically stressed conditions led to the increased DNA damage.^52^ Thus, our data strongly suggest an impaired nuclear envelope status in the *Nup133*^-/-^ neural progenitor cells. Our observations also support the notion that nucleocytoplasmic compartmentalization is disturbed under conditions in which disintegrated NPCs are present.

**Figure 7:**
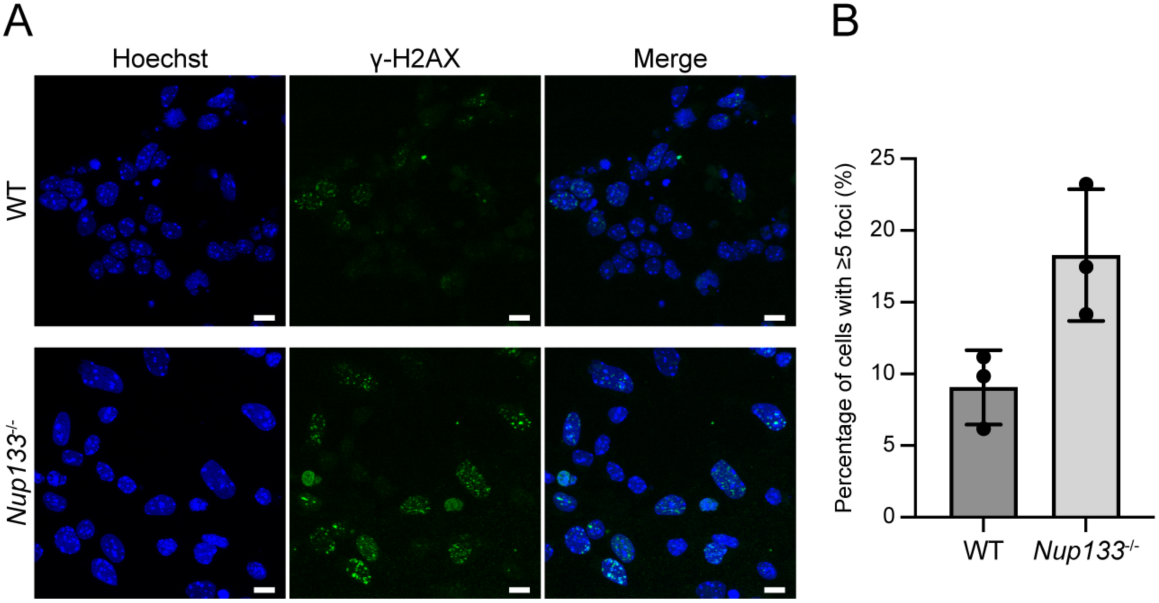
Accumulation of DNA damage is observed in the *Nup133*^-/-^ cells. (A) Representative immunofluorescence staining images of differentiating neural progenitor cells with γ-H2AX foci. Nuclei are counterstained with Hoechst 33342 and shown in blue. Scale bars, 10 μm. (B) Mean percentage of nuclei with ≥ 5 γ-H2AX foci in the wild-type and *Nup133*^-/-^ neural progenitor cells. Data are from three biological replicates, and at least 340 nuclei are analyzed for each condition. Each data point represents the percentage calculated from individual dataset. Error bars denote standard deviations.

### Over-stretching of NPCs occurs in the wild-type neural progenitor cells and human cells

Within the wild-type neural progenitor dataset, we found a subset of NPCs that exhibit abnormally large diameters, indicating that the over-stretching of NPCs could occur even without perturbation of the NPC scaffold architecture, albeit at a lower frequency. To better understand this potential over-stretching process of NPCs within the wild-type cells, we further analyzed the architectures of the over-stretched NPCs in the wild-type neural progenitor cells using TM. Similar to those in the *Nup133*^-/-^ neural progenitor cells (Fig. 6B), over-stretched NPCs in the wild-type neural progenitor cells often possess the CR at a position distant from the nuclear membranes (Fig. 8A, S7A), implying that CR detachment from the membranes may be a general feature of over-stretched NPCs. However, in contrast to over-stretched NPCs in the *Nup133*^-/-^ neural progenitor cells, where CR architectures are largely impaired (Fig. S6A), we frequently observed a complete or almost complete 8-fold symmetric CR in the wild-type dataset (Fig. 8A, S7A). Complete rings are observed mostly for the CR, while the NR of the over-stretched NPCs is more frequently fragmented (Fig. S7A), consistent with the idea that the CR has a more resilient architecture owing to the additional cytoplasm-specific components such as Nup358. Notably, the IRs in these over-stretched NPCs are incomplete and arranged in a distorted manner, indicating that their architectures are largely impaired.

**Figure 8:**
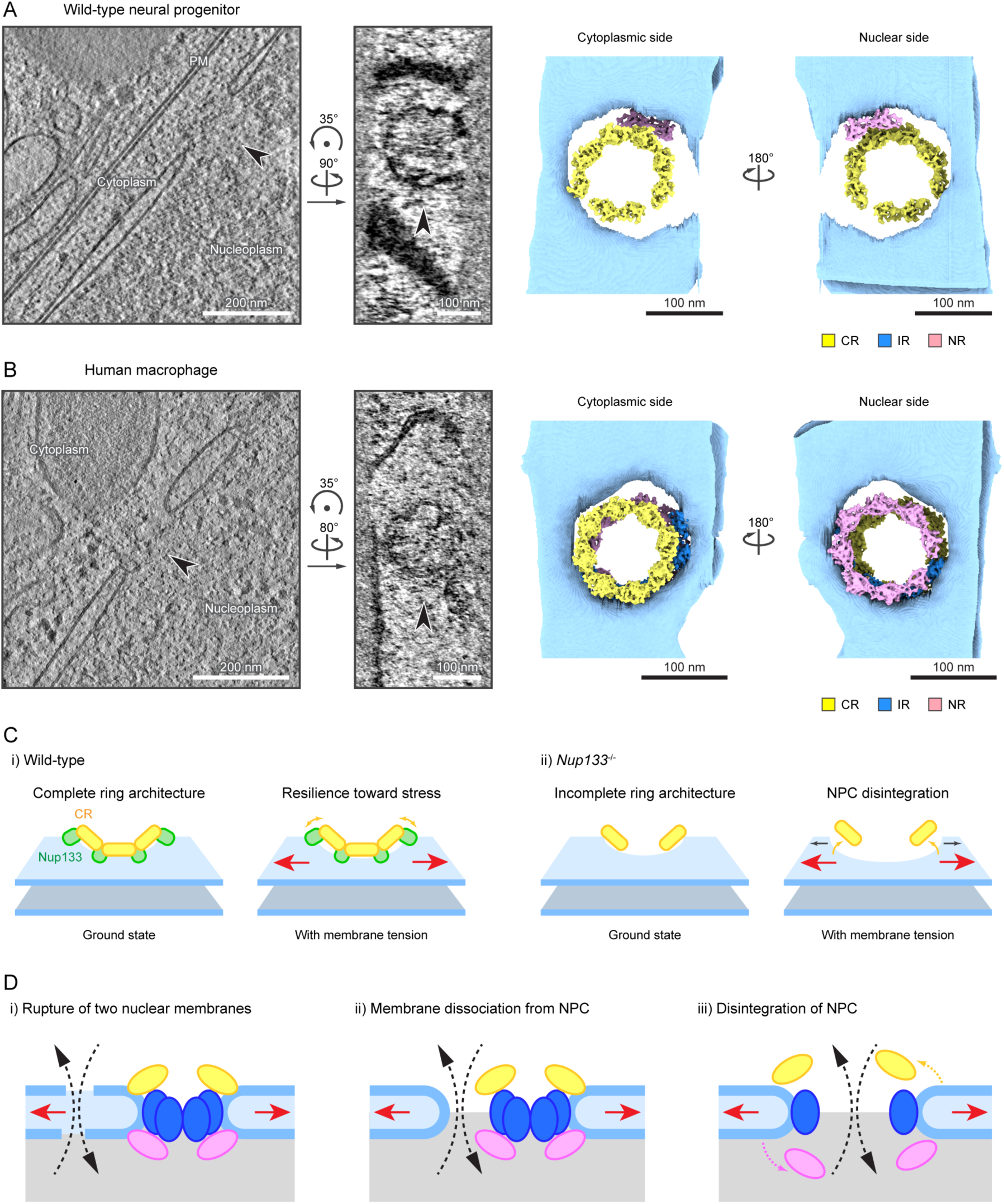
Over-stretching of NPCs can occur in wild-type neural progenitor cells and human macrophages. (A) Representative example of an over-stretched NPCs observed in the wild-type neural progenitor dataset. (B) Representative example of an over-stretched NPCs observed in human macrophage dataset. In (A) and (B), slices from the reconstructed tomograms and the pseudo-composite map generated from the results of the TM analysis are shown as in Figure 6B. Note that no IR protomer is detected for the over-stretched NPC in the wild-type neural progenitor cells shown in (A). (C) Model of NPC over-stretching in wild-type (i) and *Nup133*^-/-^ (ii) cells. In the wild-type cells, Nup133 (depicted as green spheres) mediates the inter-protomer contacts and membrane anchoring of the CR (yellow) and NR (not depicted), and thereby forms rigid ring architectures that are resilient to membrane tension (red arrows) (i). In the *Nup133*^-/-^ cells, the lack of Nup133 leads to unstable inter-protomer contacts and partial loss of CR protomers, making the CR architecture structurally less robust and more sensitive to membrane tension. Yellow arrows indicate the motion of the NPC scaffolds, and grey arrows indicate the motion of the fusion point of the outer and inner nuclear membranes. (D) Cartoons depicting possible mechanisms of nuclear envelope rupture. Nuclear envelope rupture could occur either by tearing of both the outer and inner nuclear membranes (i), by detachment of the nuclear envelope from the intact NPC scaffold architecture (ii), or by disintegration of the NPC (iii). The over-stretched NPCs observed in this study supported the scenario depicted in (iii). Black dashed arrows indicate the leakage of cytoplasmic and nuclear contents. Red arrows depict increased nuclear envelope membrane tension. The rings are colored as in Figure 6A.

Lastly, we asked if over-stretched NPCs can be found in other cell types and species. To this end, we turned to primary human macrophages, which were prepared by differentiating donor-derived monocytes.^53^ These cells are mobile and strongly adherent to the respective surfaces, implying that their nuclei are exposed to a substantial degree of mechanical stress. By acquiring cryo-electron tomograms from adherent macrophages, we indeed found a small number of NPCs with diameters of 140–150 nm in these cells (Fig. 8B, S7B). Our TM analysis revealed that these potentially over-stretched NPCs possess structural features consistent with those observed in the over-stretched murine NPCs within the wild-type neural progenitor cells, namely a membrane-detached complete CR and partially impaired NR, although some of the NRs are also detached from the membrane (Fig. 8B, S7B). Moreover, similar to the over-stretched murine NPCs, no complete IR was detected in the over-stretched NPCs from human macrophages (Fig. 8B, S7B), despite their close-to-complete CR and NR architectures, possibly implying that the IR architectures are in general more sensitive to the over-stretching than the CR and NR. Taken together, we conclude that over-stretching of the NPC architecture occurs in a similar manner in different cell types with distinct cellular states.

## Discussion

Overall, our study reveals the importance of an intact NPC scaffold architecture to safeguard the nuclear envelope during cell differentiation, where increased mechanical force is imposed on the nucleus. Moreover, our findings provide a plausible model for NPC disintegration by excess dilation under mechanical stress (Fig. 8C). In this model, the rigid ring architectures of the CR and NR of the NPC normally limit its over-stretching and disintegration to a low frequency, which may prevent large-scale damage to the nuclear envelope (Fig. 8C, panel i)). Only when the NPC is exposed to substantial stretching stress and is over-stretched to an excess degree, it starts to disintegrate, mainly by detachment of the CR from the nuclear envelope membrane. In contrast, the NPCs without Nup133 lack the stable connection between the CR and NR protomers, resulting in the formation of incomplete CR and NR. These incomplete rings fail to sufficiently restrict the NPC dilation, leading to the over-stretching and disintegration of NPCs under increased mechanical stress (Fig. 8C, panel ii)).

We propose that the NPC has to be considered as a mechanical buffer that provides additional nuclear surface area under conditions of mechanical stress. In mammalian tissue culture cells, the NPC density is about 5–10 NPCs/µm^2^,^54–56^ indicating that the NPCs constitute approximately 5–10% of the nuclear surface area. As a consequence of dilation from 90 nm to 130 nm (Fig. 1D), the area taken up by NPCs would roughly double and the nuclear surface area would increase by ∼10%. This number would be considerably larger in cell types with a higher NPC density.^57^ This expansion capacity of NPCs would allow modulation of the nuclear surface area in response to external mechanical stimuli with reduced stress imposed onto the lipid bilayer, and thereby act as a buffer for nuclear envelope stress. Such a buffering capacity could be important in dissipating mechanical stress across the nuclear envelope without impairing nucleocytoplasmic compartmentalization. With the perturbed scaffold architecture, the structural rigidity and plasticity of the NPC are lost, leading to the loss of this buffering function and higher sensitivity of the nuclei to mechanical stress.

The presence of a complete 9-fold symmetric CR architecture (Fig. 3D, E) in cells lacking Nup133 implies that the connection between two adjacent CR protomers tolerates wider angles when Nup133 is absent, further suggesting an increased flexibility of their head-to-tail contact. It is thus likely that Nup133 normally sterically limits the angles of the Y-complex head-to-tail contact and provides mechanical rigidity to these interfaces, thereby securing the overall integrity of the ring architectures. Intriguingly, this steric restriction in the head-to-tail contact may additionally play an important role for proper assembly of the NPC. Across species, the 8-fold symmetric architecture of the NPC is strictly conserved,^58^ and an NPC architecture with non-canonical symmetry has rarely been reported, with, to the best of our knowledge, only one electron microscopic study showing the presence of 9-fold and 10-fold symmetric NPCs within *Xenopus* oocyte at low frequencies.^39^ Thus, the presence of non-canonical 7-fold and 9-fold symmetric NPCs at a percentage of approximately 5% each within the *Nup133*^-/-^ mES cells is a striking phenotype, and clearly indicates a perturbed assembly due to the absence of Nup133. Since the NR is thought to serve as a primer for NPC biogenesis during both the post-mitotic and interphase NPC assembly pathways,^59^ we speculate that the architecture of the Y-complex, especially its tail structured by Nup133, determines the arrangement, stoichiometry and spacing of the assembling NRs, which subsequently regulate the geometry of the IR and ultimately of the entire NPC. The absence of fully formed NR ring architectures in the 7-fold and 9-fold symmetric NPCs, as observed in our TM analysis (Fig. 4A-E), may suggest that a complete NR ring architecture with noncanonical symmetry is initially formed at the assembling NPC, but that it disintegrates afterwards, potentially due to unstable contacts between adjacent protomers.

In addition, our TM analysis clearly revealed the presence of incomplete CRs and NRs, particularly within the *Nup133*^-/-^ cells, and thereby raised the intriguing possibility that the CR and NR protomers can be heterogeneously present within one NPC particle. This notion challenges the canonical assumption that all the symmetrically arranged protomers within NPCs are homogenously present and can thus be treated equally during averaging-based structural analyses. Indeed, the NPC structural heterogeneity we uncovered in the *Nup133*^-/-^ cells was not detectable in the initial STA averages of the 8-fold symmetric NPCs (Fig. 1E, 3H), highlighting the necessity of complementary approaches, such as the TM analysis, that do not require particle averaging. A heterogeneous presence of CR and NR protomers would be particularly relevant when the NPC architecture is perturbed experimentally, as in the *Nup133*^-/-^ cells, or deteriorated as an outcome of pathological conditions.^60^ In such conditions, it is also possible that the NPC architectures are heterogeneously affected among particles, and that only a subset of particles possess severe architectural defects, like the over-stretched NPCs within our dataset. The TM analysis could be beneficial for analyzing these minor populations of particles, as it does not require multiple homogeneous particles to average.

One further striking implication of our findings is that NPCs could be the sites where nuclear envelope rupture occurs. Prior studies have extensively investigated nuclear envelope rupture using cell biological approaches,^43,45,46,61,62^ and have established certain hallmarks and indicators of a rupture event, such as the leakage of nuclear content into the cytoplasm, the detection of cytosolic DNA by cGAS, the accumulation of nuclear DNA damage, or the detection of ESCRT-III-dependent nuclear envelope resealing. Due to the fact that nuclear envelope rupture events are rare and transient, it remains technically challenging to firmly establish that the NPC disintegration events we observed in our tomograms spatiotemporally coincide with those cell biological hallmarks. More specifically, since we are currently unable to simultaneously track the site of nuclear envelope rupture by light microscopic assays and to perform electron microscopy imaging at the very same site, we acknowledge that our experimental results provide only correlative evidence. Nevertheless, our data point to the need to revisit the prevailing hypothesis that nuclear envelope rupture occurs by the simultaneous tearing of both outer and inner nuclear envelope lipid bilayers (Fig. 8D, i)). Given the observations in our cryo-electron tomograms, two possible additional scenarios have to be considered as alternatives. In these cases, the nuclear membranes would remain unaffected and detach from the intact NPC scaffold at the fusion point of the outer and inner nuclear membranes^10^ (Fig. 8D, ii)), or the NPC scaffold would disintegrate and generate a larger membrane opening (Fig. 8D, iii)). Further in-depth investigations will be required to address this in more detail. Nonetheless, our proposed model is in line with previous findings that yeast strains lacking Y-complex Nups, including Nup133, are sensitive to DNA damaging reagents^63^ or to impaired DNA replication and repair.^64^

Multiple possibilities have been proposed as underlying mechanisms of the differentiation defect of the *Nup133*^-/-^ cells, such as impaired nuclear basket assembly^20^ or altered gene regulation.^22^ Our findings now point to an additional possibility that impaired nuclear envelope integrity, due to the disintegration of aberrant NPCs, may impact the cell differentiation process. Intriguingly, a similar neuronal differentiation defect was also observed in mES cell lines devoid of other Y-complex components Nup43 and Seh1,^21^ supporting our notion that the differentiation defect is likely attributable to impaired NPC architecture, rather than specific functions of individual Nups. The differentiation-specific impact of the Nup depletion however still remains to be fully defined. Nonetheless, we speculate that, since the size of the over-stretched NPC is larger and more variable in the differentiated *Nup133*^-/-^ cells (Fig.5D), the NPC disintegration and its damage to the nuclear envelope integrity may be exacerbated during cell differentiation. Here, the nuclear integrity gets more severely perturbed as the nucleus is exposed to increasing level of mechanical stress, and cells may fail to properly differentiate when the damage level exceeds the permissible degree. In addition to directly impairing nuclear envelope integrity, over-stretched NPCs might release the tension transmitted from the cytoskeleton networks, and thereby possibly perturb nuclear mechanosensing, which plays a key role in cell differentiation.^34^ Although the exact contribution of NPC over-stretching and disintegration to the differentiation defect awaits further investigation, the postulated requirement of an intact NPC scaffold in a physiological context is further supported by its link to human genetic diseases. Specifically, mutations in the *Nup133* gene cause steroid-resistant nephrotic syndrome (SRNS)^14^ and Galloway-Mowat syndrome,^17^ which involves neurological abnormalities in addition to the nephrotic syndrome. Intriguingly, some of the disease-causing mutations in Nup133 were demonstrated to disrupt its interaction with Nup107,^14,17^ and conversely, a causal mutation of SRNS in the *Nup107* gene is also known to disturb its interaction with Nup133.^12,14^ With these mutations, incorporation of Nup133 into the Y-complex would be perturbed, and the NPC architecture would be impaired in a mild, but likely similar manner to what we observed for the *Nup133*^-/-^ cells. Moreover, causal mutations of SRNS have also been found in two other Y-complex Nups, Nup160 and Nup85.^14^ Overall, these findings underline the physiological importance of an intact NPC scaffold architecture.

In summary, our study illustrates that the perturbation of the NPC scaffold impacts the structural completeness and stability of the NPC in a heterogenous manner, rather than causing homogeneous and large-scale structural impairment, and thereby provides insights into how mutations in Nups could cause various and complex phenotypes in cells. Moreover, this study also uncovers a critical role of the NPC scaffold in protecting membrane openings from excess expansion, whereby the NPC scaffold may be conceived as an annular spring with elastic properties that safeguards the nuclear envelope. We envision that this “safeguarding” function of the NPC should be relevant for various biological processes that impose mechanical stress on the nucleus, such as cell adhesion, differentiation or migration. The importance of the nuclear lamina for such processes has been well recognized, whereas the NPC has received less attention in these contexts. Our findings thus shed light on the previously unrecognized importance of the NPC architecture for the maintenance of nuclear integrity.

## Materials and Methods

### mES cell culture and neural progenitor differentiation

For mES cell culture, the wild-type HM1 line^65^ and the HM1-derived *Nup133^-/-^* clone (#14) described previously^20^ were used. Mouse embryonic stem cell medium was prepared by supplementing EmbryoMax® DMEM (Millipore, SLM-220-B) with 15% FBS (Gibco, 10270-106), 2 mM L-Glutamine (Gibco, 25030-024), 1 × non-essential amino acids (Gibco, 11140-050), 1 × EmbryoMax® nucleosides (Millipore, ES-008-D), 110 μM β-mercaptoethanol (Gibco, 21985-023) and 1 × penicillin/streptomycin (Gibco, 15070-063). mES cells were cultured on mitomycin-treated mouse embryonic fibroblasts (feeder cells) plated on 0.1% gelatin (Sigma-Aldrich) in mouse embryonic stem cell medium supplemented with leukemia inhibitory factor (LIF, ESGRO, Millipore, final 1000 U/ml), and kept at 37°C with 5% CO_2_. Cells were used within eight passages after thawing frozen vials to ensure the quality of the stem cell culture. The neural progenitor differentiation was performed following the previously published protocol.^38^ Briefly, after two passages on feeder cells, mES cells were transferred to gelatin-coated plates without feeder cells, and cultured for two to four passages in the presence of LIF. Subsequently, cells were trypsinized and 4 × 10^6^ cells were transferred to non-coated 10-cm Petri dishes (Greiner, 633102) and cultured for eight days within DMEM supplemented with 10% FBS (Gibco, 10270-106), 2 mM L-glutamine (Gibco, 25030-024), 1 × MEM non-essential amino acids (Gibco, 11140-035) and 100 μM β-mercaptoethanol (Gibco, 31350-010) without LIF. The culture medium was changed every two days, and after four days, 5 μM retinoic acid (Sigma, R2625) was added to the medium. After eight days of culture, the aggregates of neural progenitors were dissociated by trypsinization at 37°C for 5 min in the water bath, and the dissociated cells were either immediately used for cryo-EM grid preparation as described below, frozen within the medium supplemented with 10% DMSO and stored until use, or directly used for subsequent neuronal differentiation. For the neuronal differentiation, dissociated cells were seeded on glass bottom plates (ibidi, 80827) coated with poly-DL-ornithine (Sigma, P8638) and laminin (Roche, 11243217001) at approximately 1–2 ×10^6^ cells/cm^2^, and cultured in N2 medium (1:1 mixture of DMEM (Gibco, 21969-035) and Ham’s F-12 nutrient mix (Gibco, 21765-029), supplemented with 2 mM L-glutamine, 25 μg/ml insulin (Sigma, I6634), 50 μg/ml transferrin (Sigma, T1147), 20 nM progesterone (Sigma, P8783), 100 nM putrescine (Sigma, P5780), 30 nM sodium-selenite (Sigma, S5261), 50 μg/ml BSA (Sigma, A9418), and 1 × penicillin/streptomycin (Gibco, 15140-122)) for two days. The medium was changed to fresh N2 medium 2h and one day after plating. Images of the cells were acquired on a Nikon Ti-2 widefield microscope (Nikon), using a 10× objective.

### Cryo-EM sample preparation

For the grid preparation of mES samples, the cells were treated with double thymidine block and synchronized at the beginning of S-phase to obtain homogeneous cell population with similar nuclear size. Specifically, mES cells were cultured within mouse embryonic stem cell medium supplemented with LIF for one day after passaging, and subsequently cultured within the medium containing 2 mM thymidine (Sigma, T9250) for 18 h. Cells were then washed and cultured without thymidine for 9 h. After this short release from the cell cycle arrest, the cells were again cultured in the presence of 2 mM thymidine for 18 h. The arrested cells were trypsinized and resuspended into the fresh culture medium at the density of 2–4 × 10^6^ cells/ml for subsequent use.

Cryo-EM grid preparation was performed using a EM GP2 plunger (Leica Microsystems), with a set chamber humidity of 70% and temperature of 37°C. After trypsinization of the mES cells, 4 μl of the cell suspension was applied onto a glow-discharged holey SiO_2_ grid (R1/4, Au, 200 mesh, Quantifoil) mounted on in the plunger chamber, and 2 μl of the culture medium was applied onto the back side of the grid for the reproducible blotting. Subsequently, the grid was blotted from the back side for 10 s with Whatman filter paper, Grade 1, and plunge frozen in liquid ethane. For the neural progenitor sample, the dissociated cells from the cell aggregate formed after the eight-day culture within non-coated dishes were plunge-frozen using the same procedure, but without a double thymidine block.

Plunge-frozen samples were cryo-FIB milled using an Aquilos microscope (Thermo Scientific). More specifically, the loaded samples were first coated with a layer of organometallic platinum using a gas injection system for 10 s, and subsequently sputter-coated with inorganic platinum at 1 kV voltage and 10 mA current for 20 s. Milling was performed in a stepwise manner, with decreasing Ga ion beam current steps of 1 nA, 500 pA, 300 pA, 100 pA, and 30 pA at a stage tilt angle of 15°. Micro-expansion joints^66^ were generated using the ion beam current of 1 nA or 500 pA alongside the lamellae to prevent lamella bending. Final polishing was performed using 30 pA ion beam current, with the target lamella thickness of 180 nm. After polishing, grids were immediately recovered from the microscope and stored in liquid nitrogen until use.

### Cryo-ET data acquisition and tomogram reconstruction

Cryo-ET data acquisition was performed on a Titan Krios G2 microscope (Thermo Scientific) equipped with a BioQuantum K3 imaging filter (Gatan), operated at 300 kV in EFTEM mode with an energy slit width of 20 eV. Tilt series were acquired using SerialEM^67^ in low dose mode at a magnification of ×33,000, corresponding to a nominal pixel size of 2.682 Å, with a defocus range of -2.5 to -4.5 μm. Images were recorded as movies of 10 frames with a total dose of 2.5 e^-^/Å^2^, and motion-corrected in SerialEM on-the-fly. Tilt series acquisition started from the stage pretilt angle of 8° to compensate for the lamella angle, and images were acquired using a dose-symmetric acquisition scheme^68^ with 2° increments, up to +64° or 66° and -52° stage tilt angle, resulting in a total dose per tomogram of ∼150 e^-^/Å^2^.

Before tomographic reconstruction, the contrast transfer function (CTF) for each tilt image was estimated using gctf.^69^ Subsequently, images were filtered by cumulative electron dose,^70^ using MATLAB implementation described previously.^71^ Next, tilt images with poor quality were discarded by visual inspection. These dose-filtered and cleaned tilt series were aligned with patch-tracking in IMOD (version 4.10.9 and 4.11.5),^72^ using the 4× binned images and 9×5 patches, and reconstructed as 4× binned, SIRT-like filtered tomograms with a binned pixel size of 10.728 Å. Tomograms with sufficient visual quality and a good patch-tracking result (typically, overall residual error value below 1 pixel at 4× binning) were selected and 3D CTF-corrected using novaCTF,^73^ and further used for subtomogram averaging workflow. For the tomogram-based NPC diameter inspection, all the reconstructed tomograms were visually examined. Only tomograms with severe defect, such as overall residual error value above 3 pixels, total angular coverage less than 60°, or large amount of ice contamination, were discarded, and remaining tomograms were used for the analysis. For the visualization of tomographic slices, 4× binned 3D CTF-corrected tomograms were deconvolved in MATLAB using tom_deconv deconvolution filter^74^ to enhance the image contrast.

### Subtomogram averaging and classification

Subtomogram averaging of NPCs was performed using novaSTA^75^ as described previously,^10^ and the detailed workflows were summarized in Figure S2A. Briefly, NPC particles were manually picked using 4× binned SIRT-like filtered tomograms, and based on this information, the particles were extracted from 8× binned 3D CTF-corrected tomograms and aligned on a whole-pore level with imposed C8 symmetry using novaSTA.^75^ After initial alignment at 8× binning, the particles were again extracted from 4× binned 3D CTF-corrected tomograms and further aligned to improve the alignment precision. After obtaining the initial whole-pore average at 4× binning, the positions of asymmetric units were determined using the distance from the center of the whole pore and C8 symmetry, and subtomograms for the asymmetric units were extracted at 4× binning. Particles with the center outside of lamellae were discarded at this step. Subsequently, subtomograms were aligned and the particles with cross correlation value to the reference below 0.1 were further discarded. Due to the large number of top views, particularly in the mES datasets, structural information along the horizontal plane of NPCs was over-represented, resulting in an average with vertically elongated features along the z axis of NPCs. To obtain more isotropic averages, subtomograms from the top views of NPCs were also discarded from the dataset at this step, and remaining particles were further processed. For the subtomogram average-based NPC diameter measurement, particles from top views were kept and the entire particle set was processed in parallel. After averaging the asymmetric unit, the alignment around each ring subunit (i.e. CR, IR and NR) was further optimized using a mask covering these target areas. Subsequently, subtomograms were re-extracted from the center of individual ring subunits at 4× binning, and further aligned to obtain final averages. The masks used for the alignment were generated with Dynamo^76^ and RELION-3.1.^77^ To generate the composite maps of the whole pore, the final individual ring averages were fitted into the average of the asymmetric unit, and the summed map was assembled into an eight-fold symmetric architecture based on the coordinates used for asymmetric unit extraction.

For the *Nup133*^-/-^ mES dataset, the whole-pore particles were subjected to reference-based classification using STOPGAP,^78^ to classify the particles with 7-, 8-, and 9-fold symmetries. To generate initial references with three different symmetries, the particles of *Nup133*^-/-^ mES NPC were first manually sorted based on the arrangement of asymmetric units. Specifically, particles were initially picked from the tomograms without symmetry judgement and subjected as a whole to the whole pore averaging at 8× binning with imposed C8 symmetry. Subsequently, asymmetric units were extracted using C8 symmetry from 8× binned tomograms and aligned as described above to obtain the initial asymmetric unit average. In parallel, asymmetric units were extracted using C60 symmetry to oversample the positions on the circumference of the NPCs, and aligned using the above-mentioned asymmetric unit average as the initial reference. After 16 cycles of iterative alignments, most particles were shifted to the actual positions of the closest subunit, and these shifted positions of the oversampled particles were visually examined to assess the symmetry of each NPC. As a result, 23, 261, and 18 NPC particles out of 411 particles showed clear 7-, 8-, and 9-fold symmetric arrangement of the asymmetric units, respectively, and these particles were aligned with C7, C8, or C9 symmetry at 8× binning to obtain three initial references for the classification. With these whole-pore averages with 7-, 8-, and 9-fold symmetries as initial references, the NPC particles were subsequently subjected to reference-based classification using STOPGAP^78^ without any imposed symmetry. Among the 411 initially picked NPC particles, 11 particles at the very edge of the tomograms contained empty areas within their subtomograms, and thus were discarded from the particle set before the classification analysis. Since STOPGAP uses stochastic methods for subtomogram alignment,^78^ classification was repeated four times using the same parameters, and particles consistently classified into same classes among all four runs were selected for further processing. The particles with 8-fold symmetry were processed as described above, whereas those with 7-fold or 9-fold symmetry were only processed at 8× binning in a similar manner, due to the low particle number and the low resolution of the averages. For the *Nup133*^-/-^ neural progenitor dataset, 193 whole-pore particles were extracted and directly subjected to the reference-based classification, using the three initial references used for the classification of the *Nup133*^-/-^ mES NPCs. Due to the low number of particles classified into 7-fold symmetric class and the presence of misclassified particles, final averages for the 7-fold symmetric class lacked readily distinguishable symmetric features in some of the classification runs. Thus, to reliably separate 7-fold and 8-fold symmetric particles, classification was performed eight times, and five runs that yielded clear 7-fold and 8-fold symmetric averages were used for selecting the consistently classified particles. No particle was classified as 9-fold symmetric NPC in any of the classification runs. 157 particles consistently classified as 8-fold symmetric NPCs were further processed as described above to obtain final averages of the CR, IR and NR. The number of particles used for individual subtomogram averages are summarized in table S1.

Subtomogram average-based NPC diameter measurement was performed as described previously.^24^ For the manual measurement of the NPC diameter, 4× binned SIRT-like filtered tomograms were visually examined, and the distance between two opposite nuclear envelopes at the outer-inner nuclear envelope fusion point was measured using the distance measurement function in 3dmod.^72^ When NPCs exhibited ellipsoidal or distorted shape, the measurement was performed at the widest section. Figures were prepared using UCSF ChimeraX.^79^

### Human macrophage sample preparation and data acquisition

Monocyte-derived macrophages (MDM) were obtained from human peripheral blood mononuclear cells (PBMC) isolated from buffy coats of healthy donors as described previously.^53^ Buffy coats were obtained from anonymous blood donors at the Heidelberg University Hospital Blood Bank according to the regulations of the local ethics committee. MDM cells were cultured in RPMI 1640 medium (Thermo Fisher Scientific) supplemented with 10% heat inactivated FBS (Capricorn Scientific GmbH, Germany), 100 U/ml of penicillin, 100 μg/ml of streptomycin (Thermo Fisher Scientific) and 5% human AB serum (Capricorn Scientific GmbH, Germany). Cells were kept in a humidified incubator at 37°C with 5% CO_2_. For the grid preparation, MDM cells were detached from the growth surface by accutase (StemCell Technologies) according to the manufacturer’s instructions. 4 × 10^4^ MDM cells were seeded on glow discharged and UV-light sterilized holey carbon grids (R2/2, Au, 200 mesh, Quantifoil), which were placed in a glass-bottomed ‘microwell’ of 35-mm MatTek dish (MatTek, Ashland, MA, USA). After seeding, cells were cultured for an additional 24 h at 37 °C. MDM cells were subsequently plunge frozen in liquid ethane at -183°C using a EM GP plunger (Leica Microsystems), with a set chamber humidity of 90% and temperature of 37°C. Before plunge freezing, 3 μl of culture medium was applied onto the grids. Subsequently, the grids were blotted from the back side for 3 s with a Whatman filter paper, Grade 1, and plunge frozen. The samples were then cryo-FIB milled using an Aquilos 2 microscope (Thermo Scientific) in a similar manner as described above. In brief, loaded samples were first coated with an organometallic platinum layer using a gas injection system for 10 s, and additionally sputter-coated with inorganic platinum at 1 kV voltage and 10 mA current for 10 s. Milling was performed using AutoTEM Cryo software (version 2.4.2, Thermo Scientific) in a stepwise manner with a Ga ion beam of 30 kV voltage while reducing ion beam current from 1 nA to 50 pA. Final polishing was performed using 30 pA ion beam current, with the target lamella thickness of 200 nm.

Cryo-ET data acquisition was performed on a Titan Krios G4 microscope (Thermo Scientific) operated at 300 kV, equipped with a E-CFEG, a Falcon 4 direct electron detector (Thermo Scientific), and a Selectris X energy filter (Thermo Scientific) operated in EFTEM mode with an energy slit width of 10 eV. Tilt series were acquired using SerialEM^67^ in low dose mode at a magnification of ×53,000, corresponding to a nominal pixel size of 2.414 Å, with a defocus range of -2.0 to -4.0 μm. Tilt images were recorded as movies of 10 frames with a total dose of 2.2 e^-^/Å^2^, and motion-corrected in SerialEM on-the-fly. Tilt series acquisition started from the stage pretilt angle of -8°, and images were acquired using a dose-symmetric acquisition scheme^68^ with 2° increments up to +52°and -68° stage tilt angle, resulting in 61 projections per tilt series and a target total dose per tomogram of ∼135 e^-^/Å^2^. Tomographic reconstruction was performed as described above, with AreTomo (version 1.33)^80^ instead of IMOD used for patch-tracking.

### Three-dimensional template matching

Three-dimensional template matching (TM) was performed using STOPGAP.^78^ To reduce the amount of computation, selected whole-pore particles were extracted from 4× binned 3D CTF-corrected tomograms with a box size of 240 or 300 voxels and subjected to the analysis. For the analysis of the 8-fold symmetric NPCs, particles located within the central 616 × 1040 × ∼200-voxel volume of 4× binned tomograms (1016 × 1440 × 500- or 600-voxel total volume) were first selected to exclude particles that are close to the edge of the tomograms and thus may be partially out of the volume. From these particles, 40 and 30 particles for the wild-type and the *Nup133*^-/-^ mES datasets, respectively, were randomly selected to have the particle sets of similar size to those of the 7-fold and 9-fold symmetric NPCs. For selecting particles, random numbers were generated using MATLAB and particles with matching array numbers were selected from the list. For the analysis of the 7-fold and 9-fold symmetric NPCs, all the particles obtained from reference-based classification were used. The individual ring averages (CR, IR, or NR) of the 8-fold symmetric NPCs from the wild-type or the *Nup133*^-/-^ mES cells, aligned at 4× binning with box size of 100 voxels, were directly used as search templates. Ellipsoidal masks that cover the protein part of the averages were used as alignment masks. Low pass filtering of ∼50 Å was applied to the templates based on the resolution of final averages. Angular search was performed with 10-degree increment without any symmetry, resulting in an angle list with 15,192 different set of Euler angles. After template matching runs, the mean and standard deviation values of the cross-correlation scores within each score volume were calculated,^42^ and the cross-correlation peaks with values larger than five standard deviations from the mean were extracted using sg_tm_generate_motl.m in the STOPGAP toolbox.^78^ The extracted peaks were inspected using ArtiaX,^81^ and only the peaks with consistent particle orientations to the NPC architectures were kept. The positions of the peaks were further visually examined together with the corresponding cross-correlation score volumes, and peaks indistinguishable from background noise were discarded. For the analysis of the number of CR and NR peaks shown in Figure 3 and Figure 4, particles with less than five IR peaks were also discarded, because the overall symmetry of the NPC (7-, 8-, or 9-fold symmetry) cannot be unambiguously judged from the arrangement of the IR peaks. Subvolumes used for the TM analysis were then visually inspected, and particles that are not fully contained within the lamellae were excluded. Since the 7-fold and 9-fold symmetric NPC particles obtained after classification nevertheless include a small number of misclassified NPCs or particles with ambiguous symmetries, these particles were furthermore discarded based on the arrangement of the IR peaks. As a result, 10 out of 34 particles, 6 out of 30 particles, 12 out of 32 particles, and 11 out of 40 particles were discarded from the datasets of the *Nup133*^-/-^ mES 7-fold, 8-fold, 9-fold symmetric NPCs and the wild-type mES NPCs, respectively. For the human macrophage dataset, selected particles were extracted from 4× binned 3D CTF-corrected tomograms (9.656 Å/pix) with box size of 300 voxels. The CR, IR, and NR averages of the wild-type mES NPC were rescaled to the corresponding pixel size using relion_image_handler,^77^ together with the corresponding masks, and used as search templates. Template matching was performed as described above, and the cross-correlation peaks with the value larger than six standard deviations above the mean were extracted and analyzed. Nuclear envelopes were segmented from deconvolved tomograms^74^ using Membrain-seg^82^ with pretrained model version 10, and the results were corrected manually using Amira Software (Thermo Scientific). For the visualization of the results, MATLAB, ArtiaX^81^ and napari^83^ were used.

### Immunofluorescence staining

Cells were seeded on 8-well glass bottom chambered slides (ibidi, 80827) coated with poly-LD-ornithine and laminin as described above, and fixed at the indicated time point after seeding in 4% paraformaldehyde for 15 min. Subsequently, cells were permeabilized in 0.1% Triton in PBS for 10 min, washed three times with PBS for 5 min each, and blocked for 30 min using 3% BSA-containing PBS (PBS/BSA). After blocking, primary antibodies diluted in PBS/BSA were applied, and samples were incubated overnight at 4°C. After washing the samples with PBS, samples were incubated with secondary antibodies diluted in PBS for 2 h at room temperature. DNA was stained with 2 μM of Hoechst 33342 (Thermo Fisher Scientific, 62249) for 10 min. Images were acquired on a Stellaris 5 confocal microscope (Leica Microsystems), using a 63×/1.40 oil objective. For quantification of Pax6-positive cells, 15 z-slices were acquired with z-step of 1 μm, while 25 z-slices were acquired with z-step of 0.6 μm for the analysis of γ-H2AX foci. Typically, images were acquired from 8 to 12 randomly selected positions within one well and all the images were used for the analysis. The primary antibodies used in this study were as follows; mouse anti-Pax6 (Developmental Studies Hybridoma Bank, 61 μg/ml, dilution 1:12), mouse anti-γ-H2AX (Merck, 05-636, 1 mg/ml, dilution 1:1000). For the secondary antibodies, goat anti-mouse AlexaFluorPlus488 conjugated antibody (Thermo Fisher Scientific, A32723, 2 mg/ml, dilution 1:2000) were used.

Image analysis was performed using Fiji.^84^ For quantification of Pax6-positive cells, nuclei and Pax6-positive areas were segmented in maximum intensity projection images by auto-thresholding, and the percentage of Pax6-positive nuclei was calculated. For counting γ-H2AX foci, nuclei were segmented in maximum intensity projection images by auto-thresholding, while γ-H2AX foci were segmented using a defined threshold value, and the number of foci larger than three pixels were counted for each nucleus. For the segmentation of γ-H2AX foci, a single threshold value was used for all the images.

## Data availability

The subtomogram averages described in this study will be deposited in the Electron Microscopy Data Bank (EMDB). The tilt series data will be deposited in the Electron Microscopy Public Image Archive (EMPIAR). Light microscopy images will be deposited in the BioImage Archive (BIAD).

## Acknowledgement

We thank Anja Becker, Eva Kaindl, and Verena Pintschovius for technical support; all the members of Beck laboratory for discussion and advice. We thank Andre Schwarz from the Max Planck Institute of Brain Research for technical advice. We thank Sonja Welsch, Mark Linder, Simone Prinz and Susann Kaltwasser from the Central Electron Microscopy facility of the Max Planck Institute of Biophysics for assistance with cryo-EM sample preparation and data acquisition. We thank Özkan Yildiz, Juan F. Castillo Hernandez, Thomas Hoffmann, and the Max Planck Computing and Data Facility for computational resources. We acknowledge support from the Imaging Facility of the Max Planck Institute of Brain Research. We thank Anke-Mareil Heuser, Maria Anders-Ößwein and Vera Sonntag-Buck for preparation of the human macrophage samples; Erica Margiotta for establishing the cryo-EM sample preparation workflow at the early stage of the project.

## Author contributions

R.T., C.O., V.D. and M.B. conceived the project. R.T. performed experiments with mES and neural progenitor cells and analyzed data. C.O. and V.D. provided mES cell lines and supported R.T. with mES cell culture. J.P.K. performed cryo-FIB milling of human macrophages, and acquired and analyzed data. V.Z. prepared cryo-EM samples of human macrophages under the supervision of H.-G.K. C.E.Z. supported R.T. with cryo-EM data analysis. S.B. supported R.T. with the analysis of DNA damage foci. B.T. supported R.T. and J.P.K. with data analysis. R.T., S.B., V.D., and M.B. wrote the manuscript. V.D. and M.B. supervised the research.

---

**Supplementary Figure 1:**
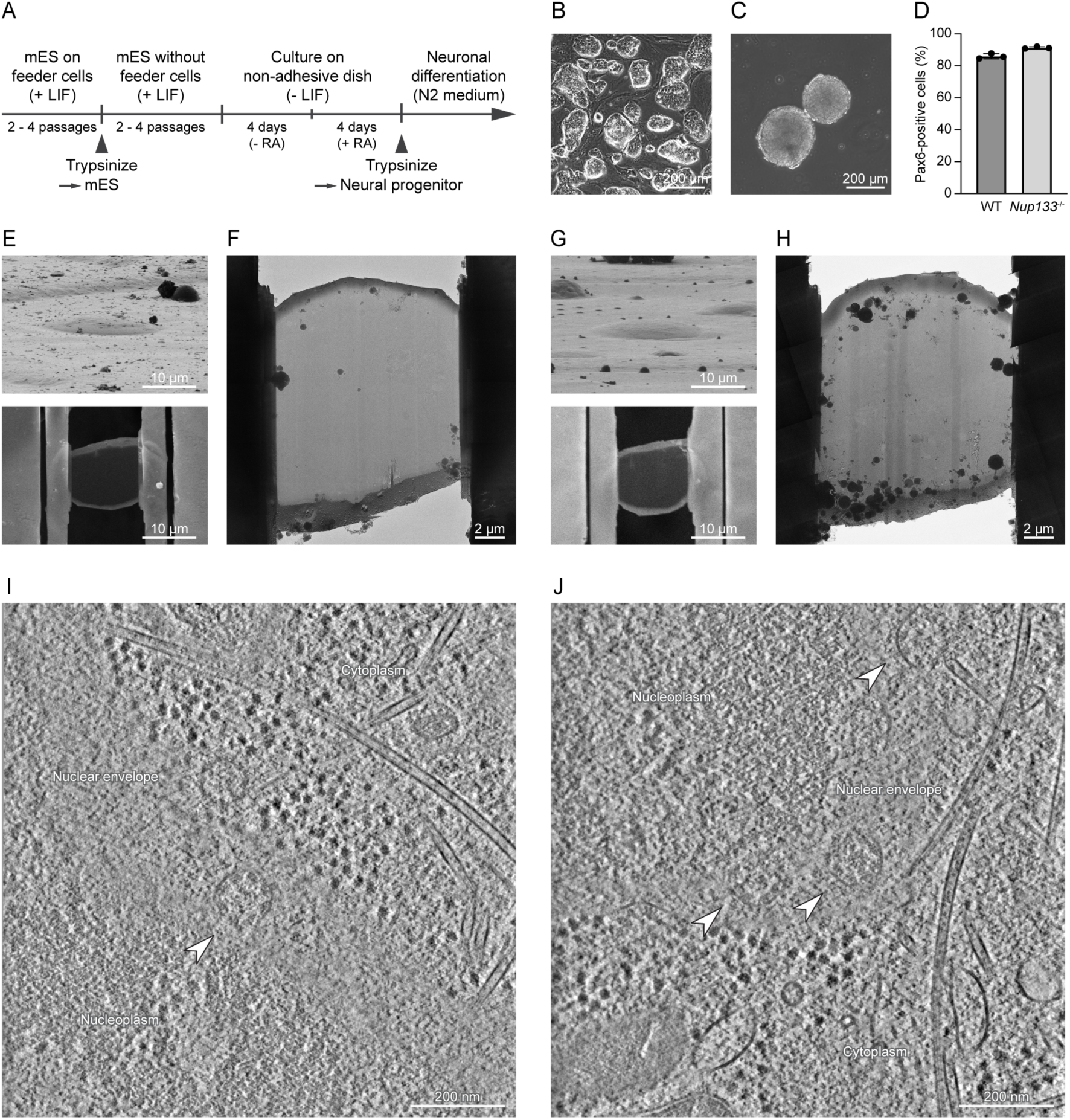
Overview of sample preparation for cryo-ET experiments. (A) Flow diagram of the neural differentiation procedure. Time-points when mES and neural progenitor cells were plunge-frozen are also indicated by arrowheads. (B) Representative image of the wild-type mES cells cultured on a layer of feeder cells. Oval colonies with bright edges are clusters of mES cells. (C) Representative image of a cluster of the wild-type neural progenitor cells. These clusters were trypsinized to obtain neural progenitor samples. (D) Quantification of Pax6-positive cells in the neural progenitor samples obtained from the wild-type and *Nup133*^-/-^ mES cells. Cells were stained 2 h after plating with Pax6 antibody. Data are from three biological replicates, and at least 300 nuclei are analyzed for each condition. Error bars denote standard deviations. (E) Representative cryo-FIB milled lamella of the wild-type mES cells. FIB view of an mES cell on a cryo-EM grid before cryo-FIB milling (top) and SEM view of the same cell after cryo-FIB milling (bottom) are shown. (F) Cryo-TEM overview of the lamella shown in (E), rotated by 180°. (G, H) Representative cryo-FIB milled lamella of the wild-type neural progenitor cells, shown as in (E) and (F). (I-J) Representative slices through reconstructed tomograms of the wild-type mES cells (I) and the wild-type neural progenitor cells (J). Top views of the NPCs are indicated by white arrowheads.

**Supplementary Figure 2:**
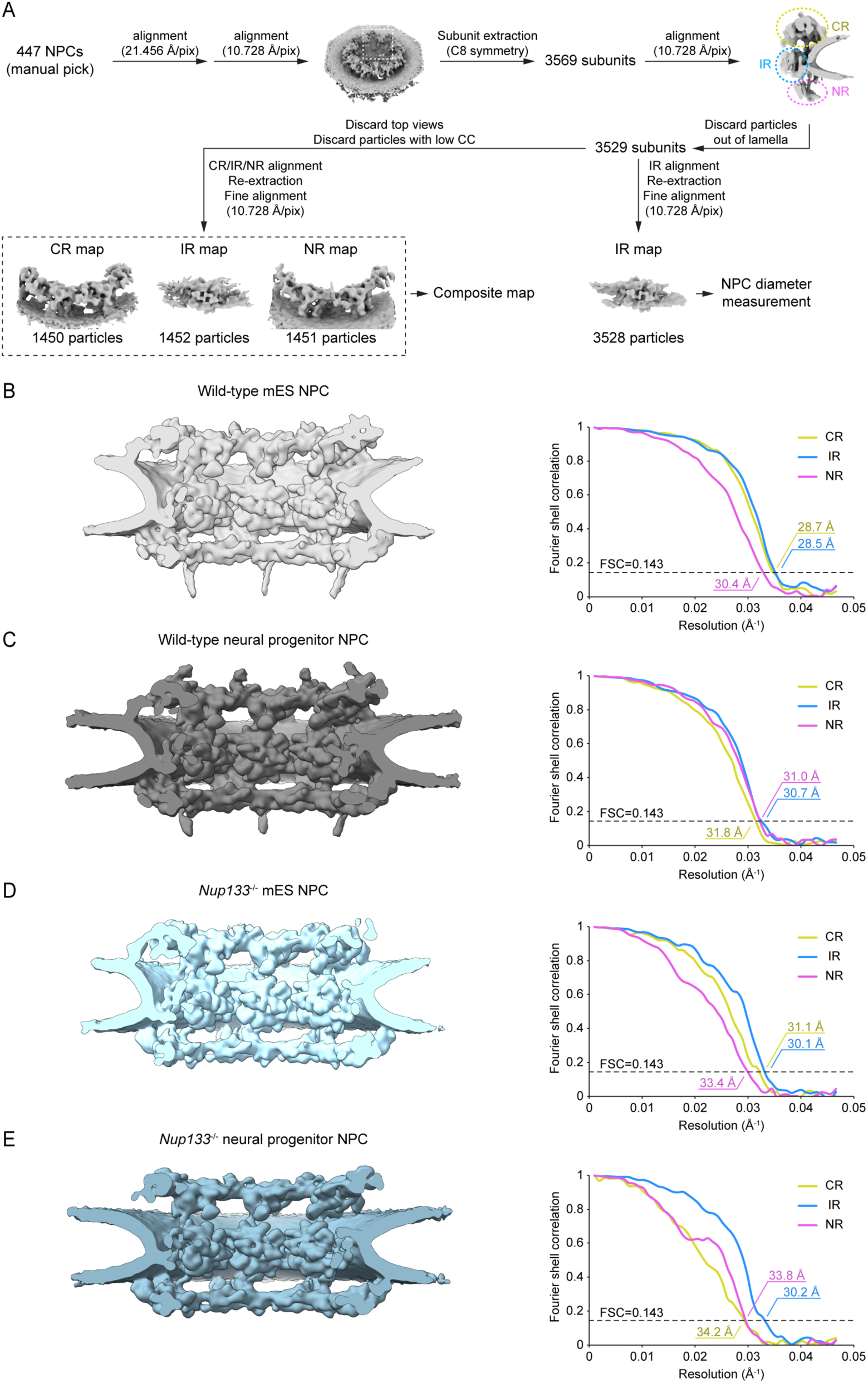
Subtomogram averages of the mES and neural progenitor NPCs. (A) The schematic data processing workflow employed for subtomogram averaging of the wild-type mES NPCs. The 8-fold symmetric NPCs from the three other datasets - *Nup133*^-/-^ mES NPC, wild-type and *Nup133*^-/-^ neural progenitor NPCs - were processed following a similar workflow. (B-E) Composite maps of the 8-fold symmetric NPCs (left) and corresponding Fourier shell correlation curves of the CR, IR and NR averages (right). The data for the wild-type mES NPC (B), the wild-type neural progenitor NPC (C), the *Nup133*^-/-^ mES NPC (D), and the *Nup133*^-/-^ neural progenitor NPC (E) are shown. Composite maps are shown as cutaway views.

**Supplementary Figure 3:**
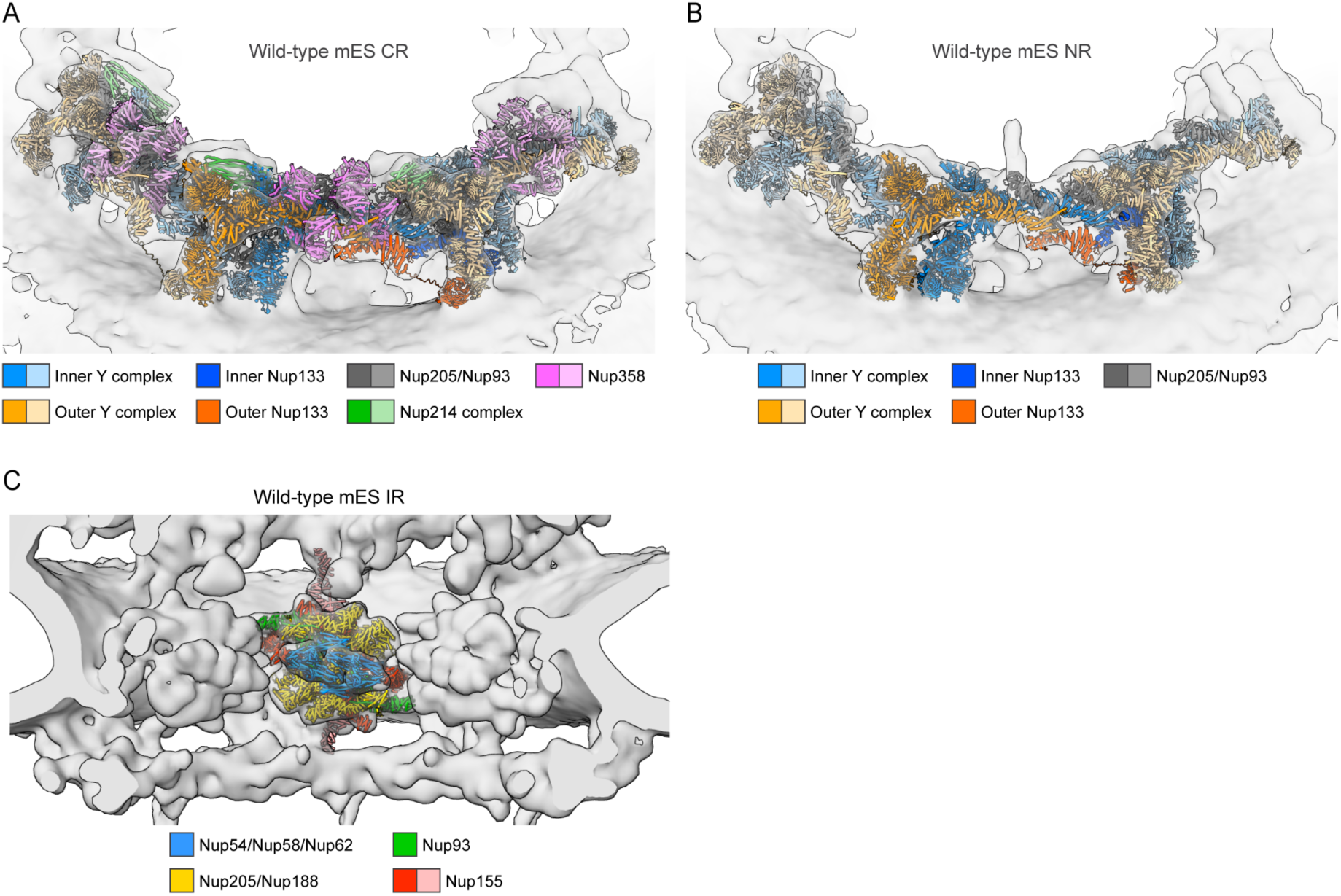
Structural comparison between murine and human NPCs. (A-C) Detailed architecture of the CR (A), NR (B), and IR(C) of the wild-type mES NPCs. The CR, NR, and IR models of the dilated human NPC (PDBID: 7R5J) are fitted into the corresponding cryo-EM maps. In (A) and (B), protomers at the center are highlighted with bold colors, while two adjacent protomers are indicated with pale colors. Note that Nup358 and Nup214 complex are CR-specific components. In (C), Nup155 molecules linking the IR with the CR or NR are colored in pale red.

**Supplementary Figure 4:**
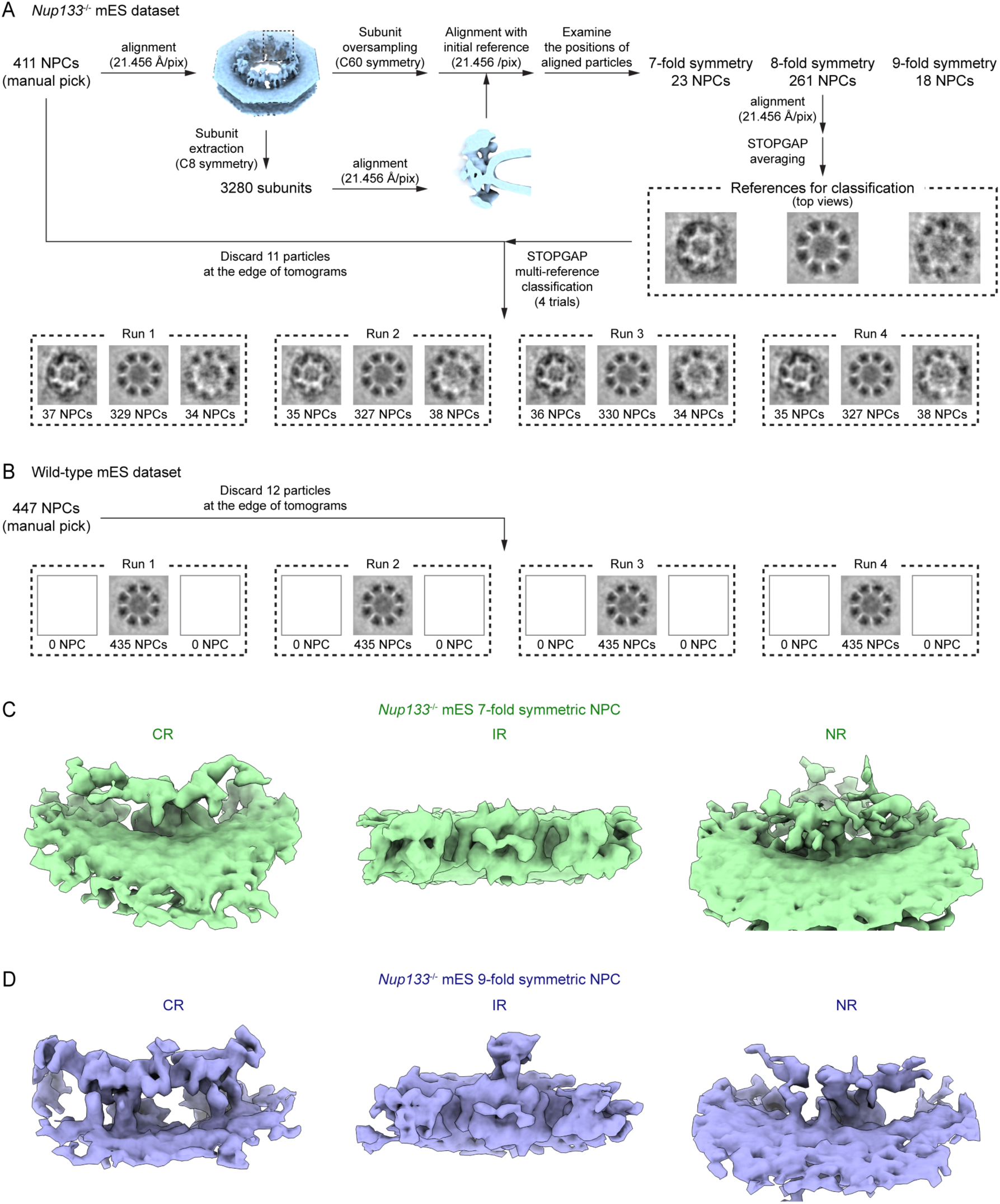
Subtomogram averaging of the 7-fold and 9-fold symmetric NPCs in the *Nup133*^-/-^ mES cells. (A) The schematic workflow of the classification of the *Nup133*^-/-^ mES NPCs. Particles with consistent class assignment among four classification runs were selected for further processing. (B) Classification of the wild-type mES NPCs. The three initial references shown in (A) were used. In contrast to the classification of the *Nup133*^-/-^ mES NPCs, all the particles were reproducibly assigned to one class with 8-fold symmetric architecture. (C-D) The cryo-EM maps of the 7-fold (C) and 9-fold (D) symmetric NPCs in the *Nup133*^-/-^ mES cells. The individual averages of the CR, IR, and NR are shown. The CR and NR averages are viewed from the same angle as in Figure 1B, and the IR average is viewed from the center of the NPC. Note that the CR and NR averages in (C) and the NR average in (D) lack interpretable structural features.

**Supplementary Figure 5:**
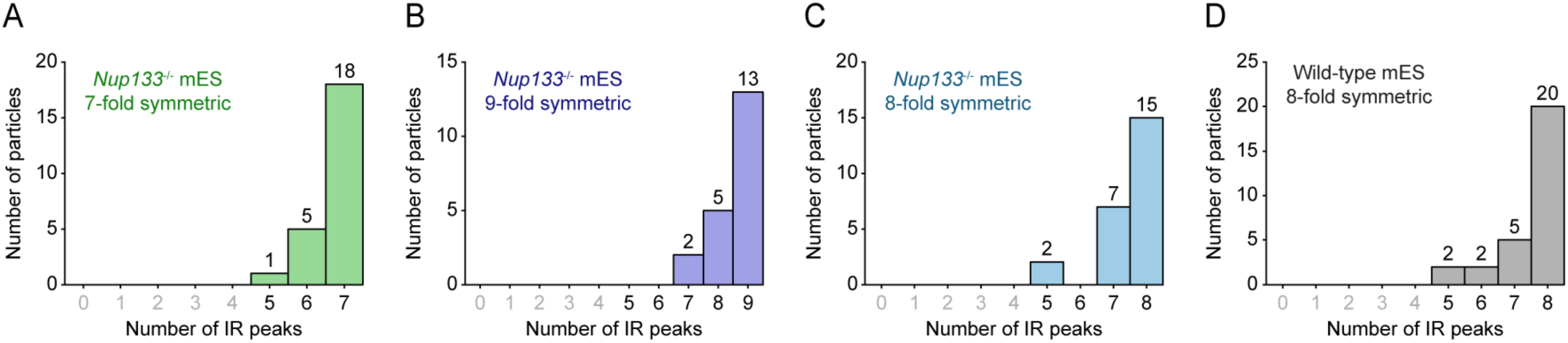
TM analysis of the mES NPCs. (A-D) Histograms showing the number of NPCs and their differing numbers of IR peaks detected by the TM analysis of the 7-fold (A), 9-fold (B), and 8-fold (C) symmetric NPCs in the Nup133-/- mES cells, and the 8-fold symmetric NPCs in the wild-type mES cells (D). Note that particles with less than five peaks were discarded from the analysis (labels colored in grey). The number of particles is indicated on top of each bar. Graphs in (A-D) correspond to the dataset presented in Figure 3C, 3E, 4B, and 4D, respectively.

**Supplementary Figure 6:**
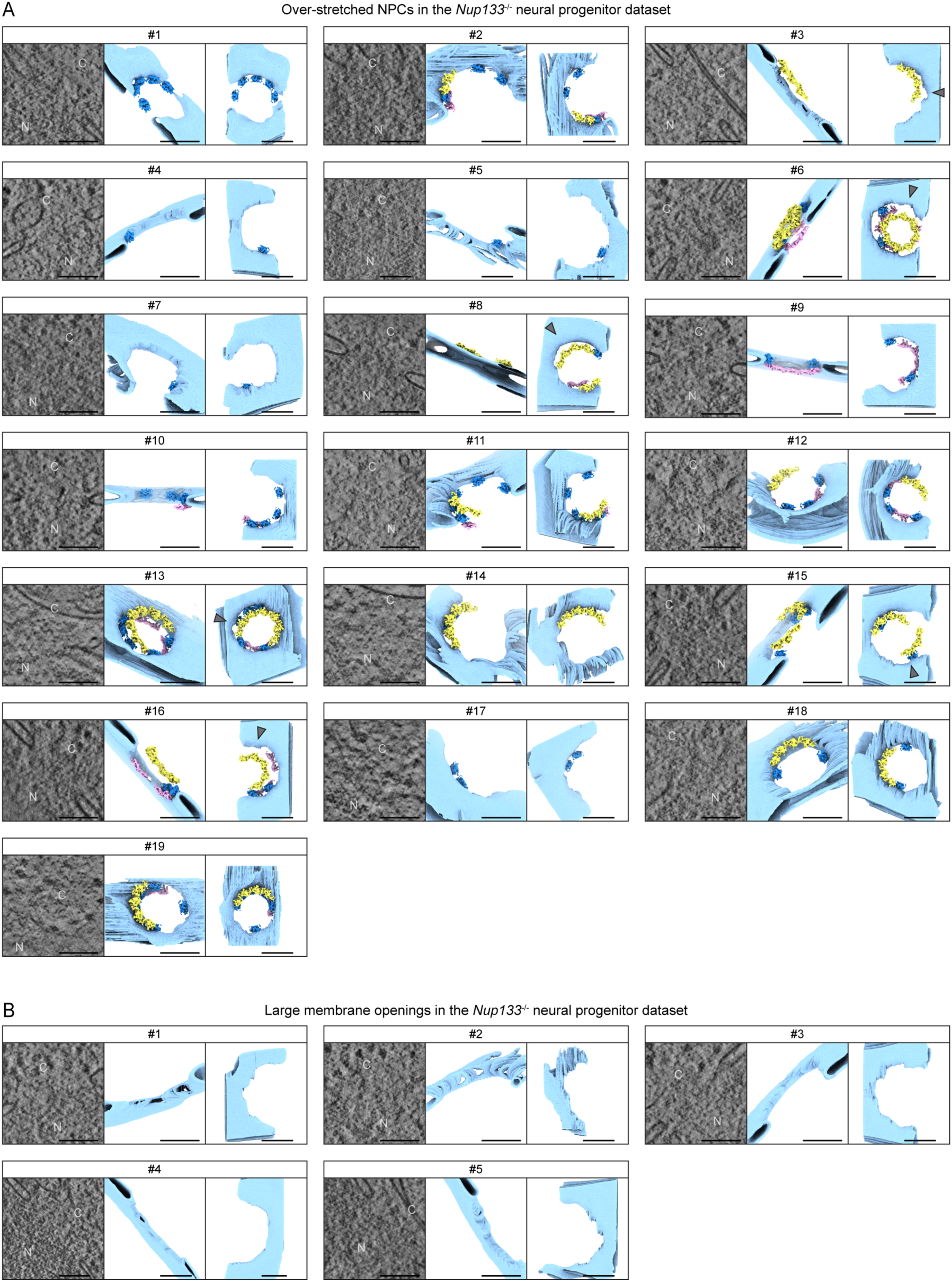
Over-stretched NPCs in the *Nup133*^-/-^ neural progenitor cells. (A) Summary of the over-stretched NPCs observed in the *Nup133*^-/-^ neural progenitor dataset. Each example of an over-stretched NPC (#1–19) includes a representative slice through the tomograms (left), segmented membranes seen from the same orientation as in the left panel (middle), and segmented membranes seen from the cytoplasmic side (right). Based on the results of the TM analysis, the CR (yellow), IR (blue) and NR (pink) maps are projected back onto the segmented membranes (middle and right panels). The CRs clearly detached from the outer nuclear membrane are highlighted with grey arrowheads. C and N in the tomographic slices indicate the cytoplasm and nucleus, respectively. The example shown in Figure 6B corresponds to #8. (B) Summary of large nuclear envelope openings observed in the *Nup133*^-/-^ neural progenitor dataset, shown in the similar manner as in (A). Note that neither the CR, IR nor NR protomer was detected in these membrane openings by the TM analysis. Scale bar, 100 nm.

**Supplementary Figure 7:**
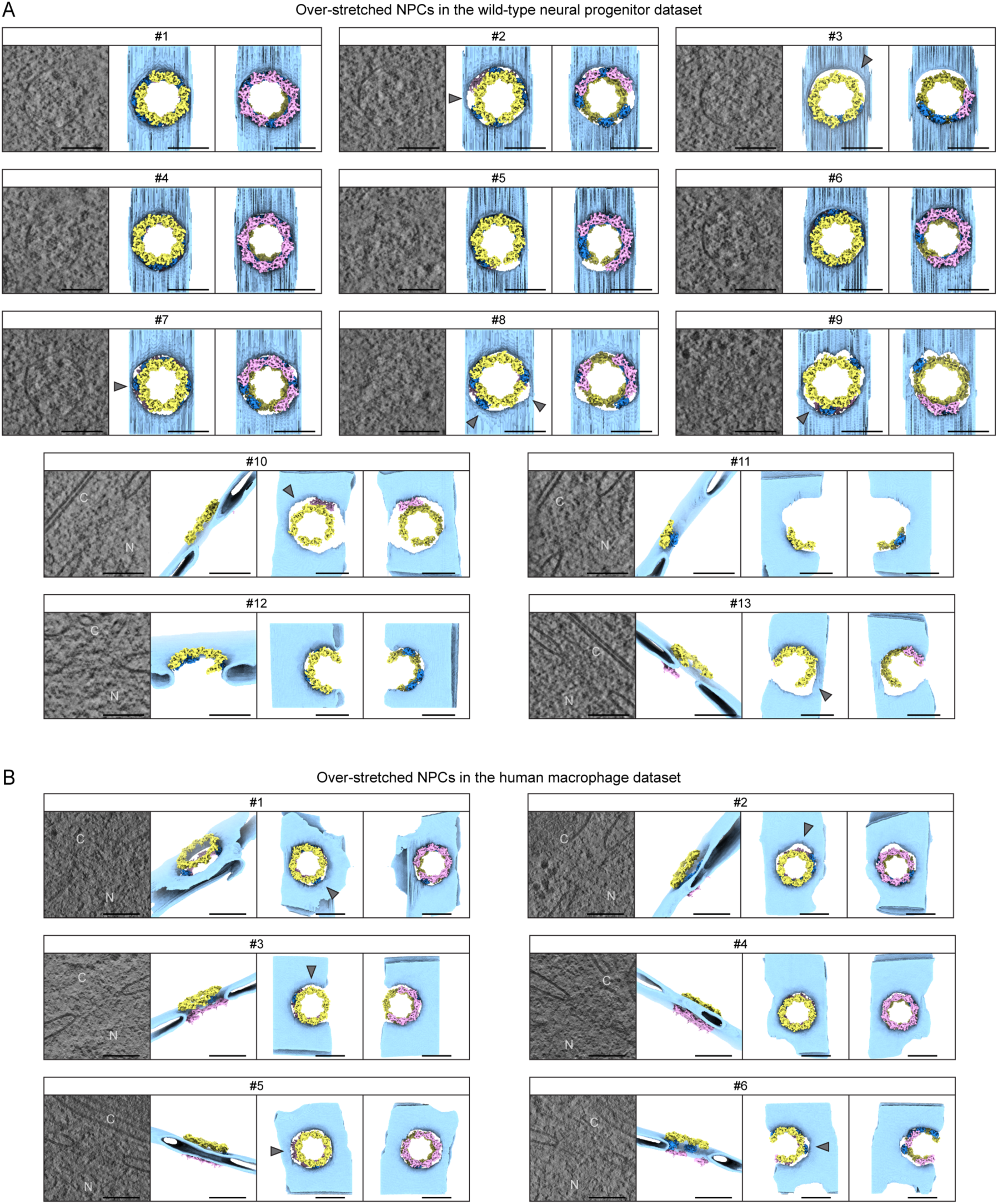
Over-stretched NPCs in the wild-type neural progenitor cells and human macrophages. (A) Summary of the over-stretched NPCs observed in the wild-type neural progenitor dataset. Each example of an over-stretched NPC (#1–13) includes a representative slice through the tomograms (left), segmented membranes seen from the same orientation as in the left panel (second from the left), and segmented membranes seen from the cytoplasmic side (second from the right) and the nuclear side (right). The CR, IR and NR maps are projected back onto the segmented membranes, based on the results of the TM analysis and colored as in Figure S6A. The CRs clearly detached from the outer nuclear membrane are highlighted with grey arrowheads. For the over-stretched NPCs oriented as top views in the tomograms (#1–9), the cytoplasmic views are omitted. C and N in the tomographic slices indicate the cytoplasm and nucleus, respectively. The example shown in Figure 8A corresponds to #10. (B) Summary of the over-stretched NPCs observed in the human macrophage dataset, shown in the similar manner as in (A). The example shown in Figure 8B corresponds to #2. Scale bar, 100 nm.

**Table S1:**
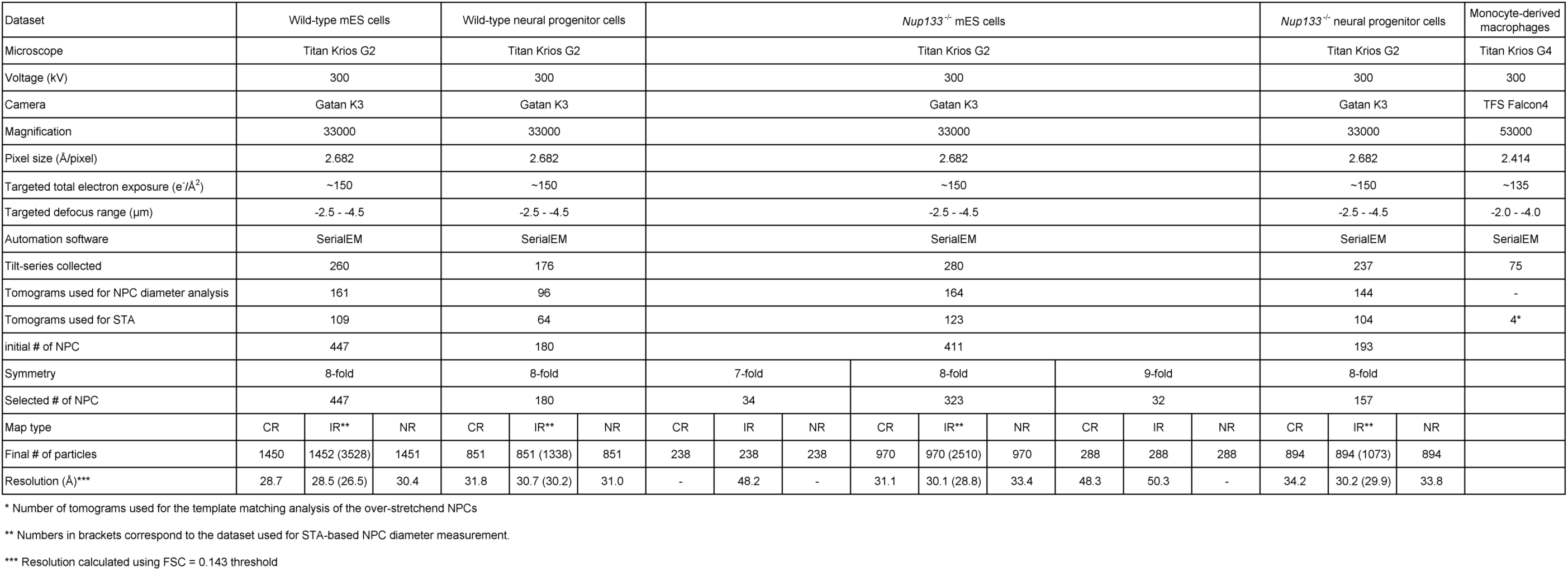
Statistics for data acquisition and subtomogram averaging.

